# Structural and functional characterization of the antigenicity of influenza A virus hemagglutinin subtype H15

**DOI:** 10.1101/2025.07.01.662631

**Authors:** Disha Bhavsar, André Nicolás León, Wei-Li Hsu, Eduard Puente-Massaguer, James A. Ferguson, Julianna Han, Patrick Wilson, Andrew B. Ward, Florian Krammer

**Affiliations:** Department of Microbiology, Icahn School of Medicine at Mount Sinai, New York, New York, USA; Center for Vaccine Research and Pandemic Preparedness (C-VaRPP), Icahn School of Medicine at Mount Sinai, New York, NY, USA; Department of Integrative Structural and Computational Biology, The Scripps Research Institute, La Jolla, CA, USA; Graduate Institute of Microbiology and Public Health, National Chung Hsing University, Taichung City, Taiwan; Drukier Institute for Children’s Health, Department of Pediatrics, Weill Cornell Medicine, New York, NY, USA; Department of Pathology, Molecular and Cell Based Medicine, Icahn School of Medicine at Mount Sinai, New York, NY, USA; Ignaz Semmelweis Institute, Interuniversity Institute for Infection Research, Medical University of Vienna, Vienna, Austria

## Abstract

Avian H15 influenza viruses are closely related to H7 viruses, but feature a unique 9-amino acid insertion in their hemagglutinin head domain, creating an additional site for antigenic variation. Here, we characterized a panel of mouse monoclonal antibodies (mAbs) raised against the A/wedge-tailed shearwater/Western Australia/2576/1979 ancestral strain, and a human mAb isolated from an H7N9 vaccinee. We found differences in binding and neutralization profiles against the ancestral strain and drifted strains of H15 isolated after 2008. MAbs that have hemagglutination inhibition activity against the ancestral strain do not show binding to drifted strains, hinting at antigenic differences in the receptor binding site. We show that the mAbs protect *in vivo* and elucidate mAb-antigen interactions using negative stain and cryo-electron microscopy. The characterization of H15 antigenicity and mechanisms of antibody-mediated neutralization expands our knowledge of this rare avian influenza virus and informs our understanding of immune pressures on viral surface glycoproteins.

## INTRODUCTION

Global climate change, habitat destruction and disruption, light pollution, and other human activities have caused physiological and behavioral changes in migratory birds (Carey, 2009; Dezfuli et al., 2022; Horton et al., 2023; Shipley et al., 2022; Wang et al., 2022; Zimova et al., 2021). These changes have often led to population and health declines as well as the intermingling of species that were formerly geographically distinct, creating the preconditions necessary for devastating viral incursions (Chen & Khanna, 2024). The dangers associated with such instances is highlighted by recent reports of H5N1 influenza A virus (IAV) outbreaks in brown skuas and gentoo penguins in South Georgia and the Falkland islands—wild bird populations in which H5N1 had not been detected previously (Banyard et al., 2024). While there is an extensive literature on seasonal and pandemic IAVs that circulate in human populations and avian IAVs with high pandemic potential, it is important to note that wild birds are also threatened by IAV pandemics, particularly when managing the fitness impacts of habitat disruption, dietary disruption, and changes in global weather patterns that generally outpace the speed of evolutionary adaptation (Boon et al., 2007; Hanmer et al., 2022; Kleyheeg et al., 2017; Sehgal, 2010). Thus, studies of rare wild bird IAV subtypes function as both proactive mitigation of potential human spillover events, which are expected to increase due to climate change, and extensions of an ethos of wildlife stewardship in the wake of manmade impacts on wild bird populations.

Hemagglutinin (HA) is a type-I viral fusion glycoprotein expressed as a homotrimer on the surface of influenza virions. Most HAs recognize sialic acid glycans as their host receptors and host specificity is thereby defined by the ability to bind moieties of α2,3-sialic acid (predominant in avian hosts) or α2,6-sialic acid (predominant in mammalian hosts)⍰⍰Nineteen influenza A virus HA subtypes(Karakus et al., 2024)⍰ have been identified in the animal reservoir of which H1, H2 and H3 have caused human pandemics so far(Krammer et al., 2018)⍰. A number of other virus subtypes—like H4, H5, H6, H7, H9 and H10—have caused zoonotic infections,(Claas et al., 1998; Gao et al., 2013; Huang et al., 2015; Kayali et al., 2011; Shi et al., 2013; To et al., 2014; Yuen et al., 1998)⍰. One of the rarest subtypes is the H15 HA. H15 is a group 2 influenza A⍰H7 and H10 HA by an ⍰ in its globular head domain, a feature not found in any other influenza virus HA subtype so far(Tzarum et al., 2017; Yang et al., 2020)⍰. H15N9 was first isolated in 1979 in Australia from a shearwater and a shelduck with additional isolations of H15N2, H15N8, and H15N9 in Australia in 1983 from ducks, terns, shelducks(Röhm et al., 1996; Sivay et al., 2013)⍰. A report from 1996 describes it as a new subtype(Röhm et al., 1996)⍰. After a long hiatus, H15N4 was found in 2008 in a teal in Russia(Sivay et al., 2013)⍰ followed by detections of H15N7 in (Sivay et al., 2013)⍰, 2012 in Sweden (H15N5 in mallard ducks) and finally in 2015 in ducks (H15N9) in Bangladesh(El-Shesheny et al., 2018)⍰ (Figure 1A). H15 has not been detected in the Americas so far and it is unclear if its main reservoir are *Charadriiformes*, *Procellariiformes,* or *Anseriformes* and if this has changed over time. So far, no human infections have been identified and H15 antibody cross-reactivity (e.g. from H3N2 infections) in human serum is among the lowest of all subtypes— (Nachbagauer et al., 2017)⍰.

**Figure 1:**
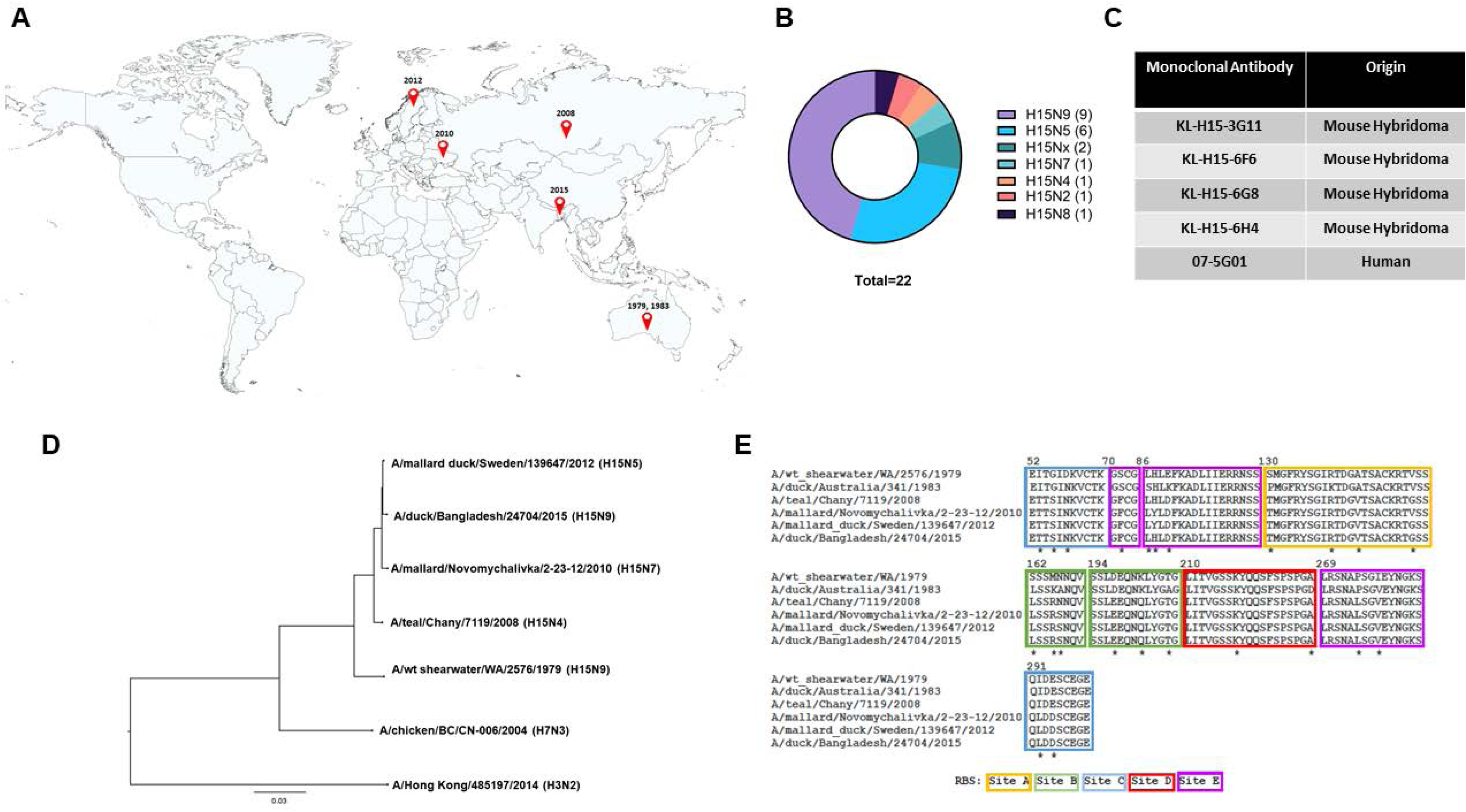
Background to H15. (A) Global map showing the locations and the years during which H15 virus strains were reported. (B) Proportion of different virus subtypes with H15 HA. Sequences of influenza virus strains with H15 HA subtype were obtained from Global Initiative on Sharing All Influenza Data (GISAID) and classified based on the NA subtype. (C) mAbs used in the study along with the source from which they were isolated. 07-5G01 was isolated from a human subject vaccinated against H7N9 virus (Henry Dunand et al., 2016), whereas the other mAbs were obtained from murine hybridomas generated by immunization with H15N5 virus. (D) The phylogenic tree representing the antigenic distance between different H15 HAs used in the study. The scale bar represents a 3% amino acid difference in sequence identity. The sequences were obtained from GenBank, and the phylogenetic tree was generated using Clustal Omega and visualized using FigTree. (E) Alignment of H15 sequences showing major antigenic sites. The amino acid sequences obtained from GenBank were aligned using Clustal Omega, and the sequences corresponding to the major antigenic sites are represented. The amino acid differences are marked with asterisks.

The Global Initiative on Sharing All Influenza Data (GISAID) only contains data for 22 H15Nx viruses (Figure 1B); given the increasing number of avian influenza sero-surveillance studies, this suggests that H15Nx viruses are rare amongst bird populations studied thus far (Hassan et al., 2020; Khare et al., 2021). There are conflicting reports on the pathogenicity of H15Nx viruses in mice and human cell lines. In one study, H15N9 failed to cause mortality or significant weight loss in mice while in another H15N1 was found to cause severe viral pneumonia and high mortality (El-Shesheny et al., 2018; Qi et al., 2014). Thus, it is likely that the HA-NA pairing as well as genotype and accessory non-structural proteins play a large role in determining the pathogenicity of H15-expressing viruses. Given the potential for IAVs to undergo genome reassortment during viral replication, the presence of mammalian virulence factors in certain H15Nx viruses emphasizes the value of proactive study to enhance pandemic preparedness. Here, we set out to test the antigenicity of H15 viruses using monoclonal antibodies (mAbs) raised against the H15 strain from 1979 as well as polyclonal sera raised in mice. We further characterized the identified mAbs regarding their mechanism of action, protective effect, and epitopes. Collectively, this study provides the first molecular characterization of H15 HA antigenicity, reports the first structures of mAbs against this rare subtype, and highlights the need for further work on antigenic profiling of rare avian IAV glycoproteins both in mammals and in their native hosts.

## RESULTS

### Monoclonal antibody generation

Previously, human mAb 07-5G01 was isolated from an individual who received two doses of a live-attenuated cold-adapted influenza A/Anhui/1/2013 (H7N9) vaccine followed by an inactivated virus vaccine boost 12 weeks later (Henry Dunand et al., 2016). Interestingly, mAb 07-5G01 exhibited neutralizing activity against the A/wedge-tailed shearwater/Western Australia/2576/1979 (A/wt shearwater/WA/2579/1979) (H15N9) virus and bound to recombinant H15 HA.

H15 reactive mouse mAbs were generated using a hybridoma based approach. Female BALB/c mice were infected with a sublethal dose of the A/wt shearwater/WA/1979 (H15N5, A/PR/8/34 reassortant) virus and subsequently boosted with an inactivated preparation of the same virus. Splenocytes of these mice were fused with the SP2/0 myeloma cells to generate hybridoma cells. The supernatants from these cells were screened for binding to A/wt shearwater/WA/1979 recombinant H15 in an enzyme linked immunosorbent assay (ELISA). Four antibodies, KL-H15-3G11 (IgG_2a_), KL-H15-6F6 (IgG_2a_), KL-H15-6G8 (IgG_2a_) and KL-H15-6H4 (IgG_2b_) were found to bind the recombinant H15 and were selected for further characterization (Figure 1C).

### The isolated mAbs show limited reactivity to a panel of H15 hemagglutinins

To determine the breadth of antigenic activity of the mAbs against a range of H15s, the full length amino acid sequences of selected H15 sequences were aligned using Clustal Omega and analyzed phylogenetically via neighbor-joining tree method (Figure 1D). The sequence alignment revealed mutations in the putative major antigenic sites of the H15s, some of which are consistent across the strains reported since 2008 (Figure 1E). ELISAs were performed against a panel of recombinant H15 proteins from viruses spanning from 1979 to 2015, when the most recent H15 virus isolate was reported. While all five mAbs bound to the recombinant H15 from the A/wt shearwater/WA/1979 strain, only KL-H15-6H4 showed binding against the recombinant H15 from the strains reported since 2008 (Figure 2A). All mAbs showed binding in ELISA against recombinant chimeric 15/3 hemagglutinin (cH15/3) which consists of head domain from A/wt shearwater/WA/1979 H15 and stalk domain from A/Hong Kong/4801/2014 H3 (Chen et al., 2016), indicating that all the mAbs potentially bind the head domain of H15 HA.

**Figure 2:**
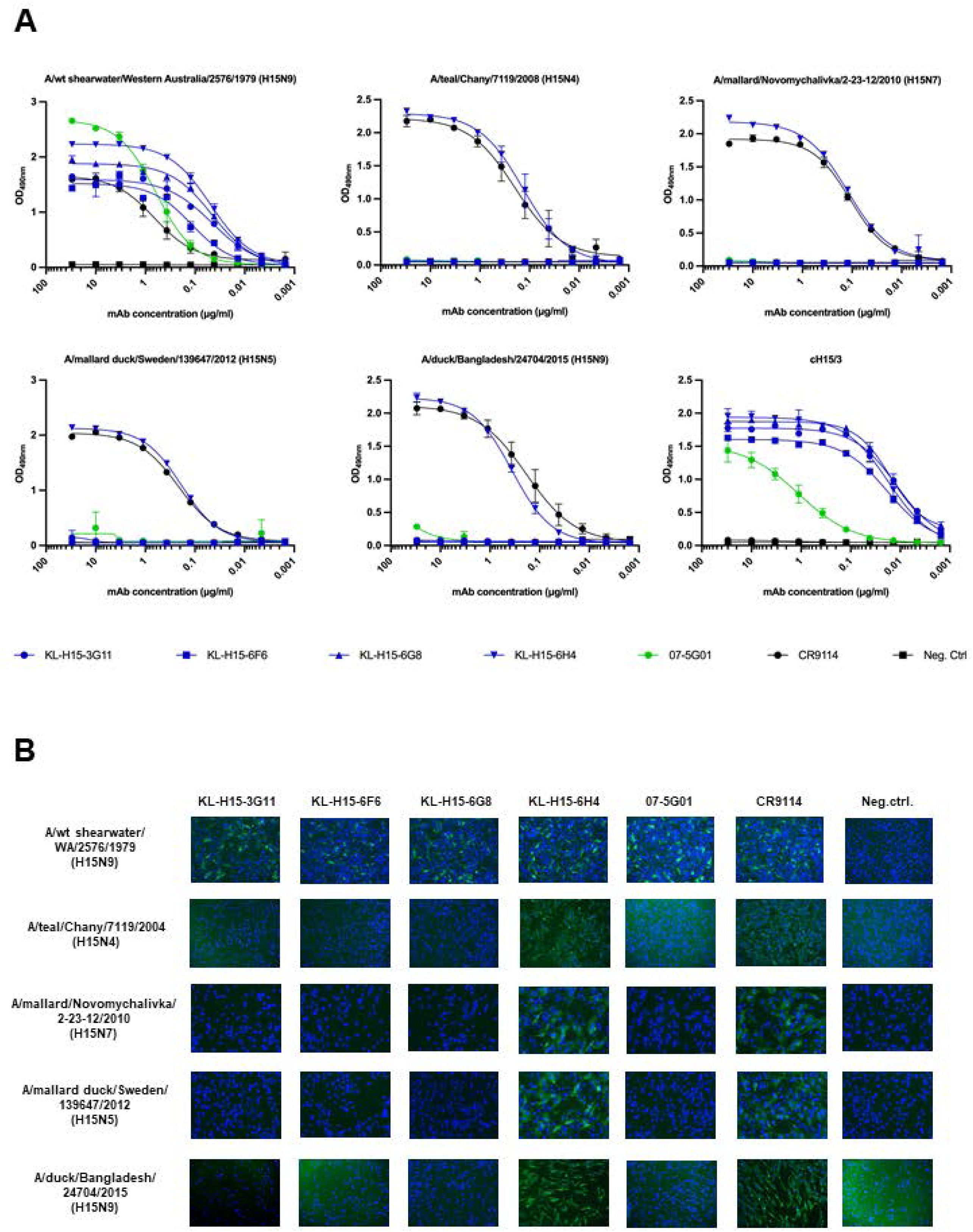
Binding of mAb panel to different H15 HAs. (A) Binding profile of mAbs to recombinant H15s from 1979 to 2015 and cH15/3 proteins as assessed by ELISA. The broadly cross-reactive HA antibody CR9114 (Dreyfus et al., 2012) was used as a positive control and anti-H6 antibody KL-H6-8H9 was used as negative control. (B) Binding of mAbs to cells expressing H15 HA. Binding of mAbs to MDCK cells infected with different H15 viruses and thus, expressing H15 on cell surface, was assessed by immunofluorescence. CR9114 (Dreyfus et al., 2012) was used as a positive control and anti-H6 antibody KL-H6-8H9 was used as negative control.

An immunofluorescence (IF) assay was also performed to test the binding of the mAbs to H15 HAs expressed on the surface of infected cells in their native conformation. Reassortant viruses (6:2) were generated with HA and NA from the A/wt shearwater/WA/1979 (H15N9), A/teal/Chany/7119/2008 (H15N4), A/mallard duck/Sweden/139647/2012 (H15N5), and A/duck/Bangladesh/24704/2015 (H15N9) strains and the internal proteins from the A/PR/8/34 (H1N1) strain. A 7:1 reassortant virus was generated for the A/mallard/Novomychalivka/2-23-12/2010 (H15N7) strain where NA from A/PR/8/34 was used. Madin Darby canine kidney (MDCK) cells were infected with the five different H15 virus strains and used for an IF assay. All mAbs showed high binding intensity to cells infected with A/wt shearwater/WA/1979, while only KL-H15-6H4 showed binding to cells infected with H15 strains isolated 2008 onwards, corroborating the results observed in ELISA (Figure 2B). Similar results were observed in ELISA against purified A/wt shearwater/WA/1979 H15N9 virus (Figure S1A).

Binding of these mAbs against A/wt shearwater/WA/1979 recombinant H15 was also tested via Western blot following reducing denaturing sodium dodecyl sulfate polyacrylamide gel electrophoresis (SDS-PAGE). All mAbs bound to H15 indicated by the presence of bands on the Western blot, although 07-5G01 seemed to bind to a lesser extent as indicated by the relatively low intensity of the band (Figure S1B).

### The majority of the isolated mAbs have neutralizing activity against the A/wt shearwater/WA/1979 strain *in vitro*

To assess whether the mAbs can neutralize the H15 viruses *in vitro*, microneutralization assays were performed. Even though the virus used to immunize the mice to generate the hybridomas had a mismatched NA (A/wt shearwater/WA/1979 H15N5), all mAbs were able to neutralize the A/wt shearwater/WA/1979 (H15N9) virus with minimal neutralizing concentrations ranging from 1 µg/ml to 0.1 µg/ml (Figure 3A). Only KL-H15-6H4 showed neutralization activity against A/teal/Chany/7119/2008 (H15N4), A/mallard/Novomychalivka/2-23-12/2010 (H15N7), and A/mallard duck/Sweden/139647/2012 (H15N5) viruses. None of the mAbs showed detectable neutralization activity against A/duck/Bangladesh/24704/2015 (H15N9) virus at the highest tested concentration.

**Figure 3:**
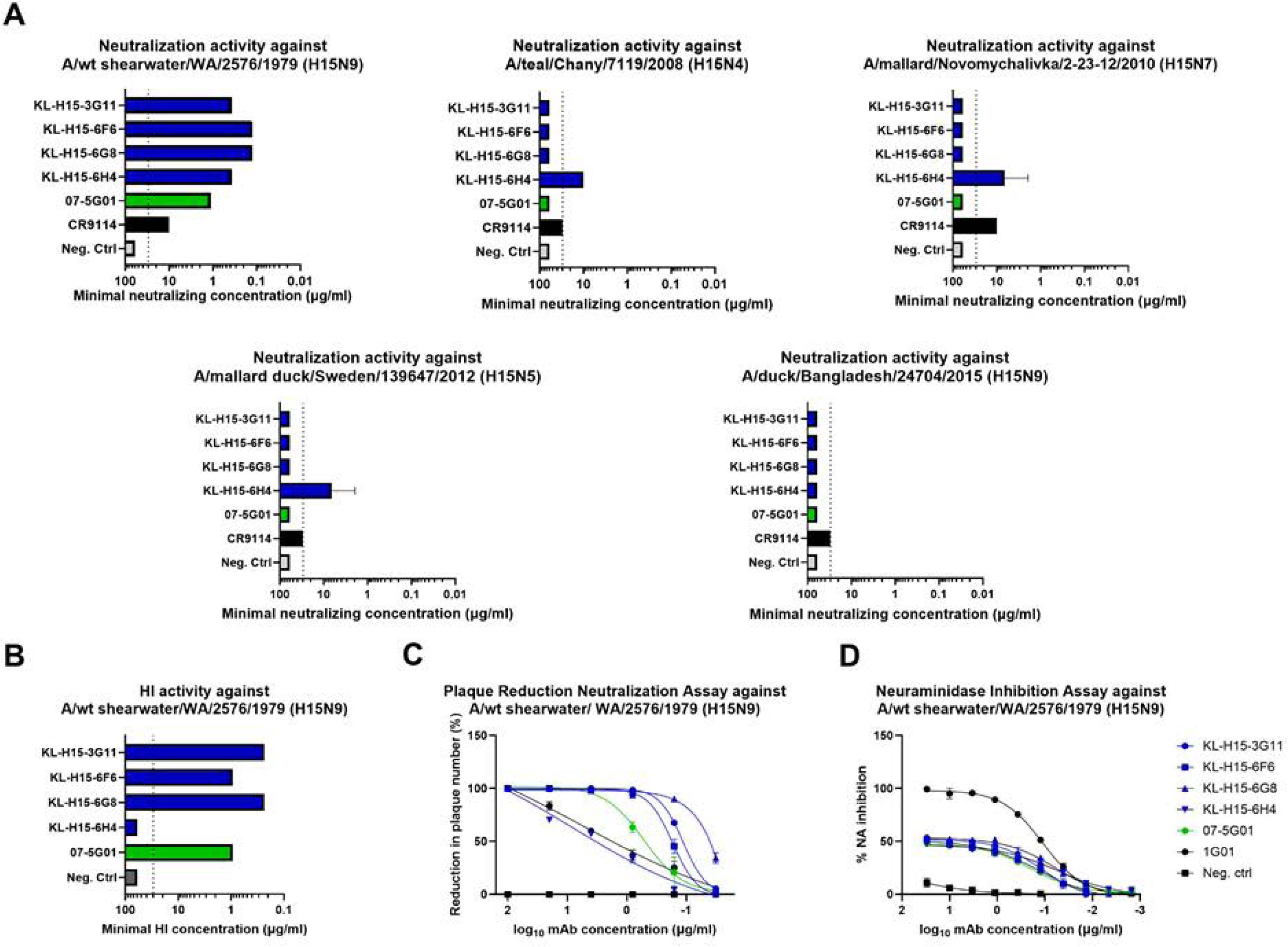
In vitro neutralization of H15 viruses by mAbs. (A) Neutralizing activity of mAbs against different H15 viruses measured by microneutralization assay. Shown here are the minimum concentrations of mAbs at which virus neutralization was observed. Broadly neutralizing HA antibody CR9114 (Dreyfus et al., 2012) was used as a positive control and anti-H6 antibody KL-H6-8H9 was used as a negative control. (B) HI activity of mAbs against A/wt shearwater/WA/2576/1979 virus. The ability of the mAbs to neutralize the virus via blocking the receptor binding site of H15 was assessed by HI assay and the minimum concentration at which HI activity was observed is shown here. Anti-H6 antibody KL-H6-8H9 was used as negative control. (C) Neutralizing activity of mAbs against A/wt shearwater/WA/2576/1979 virus measured by PRNA. Percentage reduction in number of plaques for each mAb, in five-fold dilutions starting at 100ug/ml, is shown here normalized to negative control mAb KL-H6-8H9. (D) NAI activity of mAbs against A/wt shearwater/WA/2576/1979 virus. Neuraminidase inhibition activity of the mAbs was measured by ELLA. Virus only and no-virus controls were used to normalize 0% and 100% neuraminidase inhibition activity, broadly cross reactive influenza virus neuraminidase antibody 1G01 (Stadlbauer et al., 2019) was used as a positive control, while anti-H6 antibody KL-H6-8H9 was used as negative control.

To determine if the neutralization activity of these mAbs is driven by blocking of the receptor binding site (RBS) of H15, a hemagglutination inhibition (HI) assay was used (Brandenburg et al., 2013). KL-H15-3G11, KL-H15-6F6, KL-H15-6G8 and 07-5G01 showed HI activity against A/wt shearwater/WA/1979 (H15N9) at minimum HI concentrations ranging from 0.94 µg/ml to 0.23 µg/ml (Figure 3B). While KL-H15-6H4 was observed to have neutralization activity, no HI activity was detected for this mAb, hinting at neutralization by mechanisms other than blocking of the RBS. No HI activity was observed for these mAbs against the virus strains isolated since 2008 (Figure S1C).

In addition to microneutralization assays, the neutralization activity of these mAbs was also tested in a plaque reduction neutralization assay (PRNA) against A/wt shearwater/WA/1979 (H15N9) virus. Neutralization capacity was determined by the percentage reduction in number of plaques formed by the virus incubated with dilutions of the corresponding mAb compared to an irrelevant IgG. All mAbs showed 100% reduction in plaque number at the highest tested concentration of 100 µg/ml. The 50% inhibitory concentration (IC_50_) values were 0.11 µg/ml for KL-H15-3G11, 0.17 µg/ml for KL-H15-6F6, 0.001 µg/ml for KL-H15-6G8, 8.039 µg/ml for KL-H15-6H4 and 0.52 µg/ml for 07-5G01 (Figure 3C).

Interestingly, reduction in plaque size was also observed in the wells treated with neutralizing mAbs in addition to reduction in number of plaques (Figure S2A). HA mAbs have previously been shown to inhibit neuraminidase activity and contribute to influenza virus neutralization due to steric hindrance (Rajendran et al., 2017; Wohlbold et al., 2016). To assess this, neuraminidase inhibition (NAI) assays were performed using an enzyme-linked lectin assay (ELLA). MAbs were pre-incubated with A/wt shearwater/WA/1979 (H15N9) and transferred onto fetuin coated plates. Cleavage of sialic acid was measured by binding of peanut agglutinin. No virus and no mAb controls were used to normalize 100% and 0% neuraminidase NA inhibition. Mab 1G01, a broadly cross-reactive anti-influenza virus neuraminidase antibody was used as a positive control (Stadlbauer et al., 2019). At the maximum tested concentration of 30 µg/ml, the mAbs appeared to show 50% NA inhibition (Figure 3D). KL-H15-6H4 was also able to inhibit the neuraminidase NA activity of H15 viruses isolated 2008 onwards (Figure S2B). Additionally, all mAbs except 07-5G01 were also able to induce varying degrees of detectable antibody-dependent cellular cytotoxicity (ADCC) activity when measured using a commercial reporter assay (Figure S2C).

### The mAbs confer protection *in vivo* against lethal H15 virus challenge and inhibit viral replication in mouse lungs

*In vivo* protective efficacy of the mAbs was tested in a mouse model against lethal challenge with A/wt shearwater/WA/1979 (H15N5) virus in a prophylactic setting. MAbs were administered intraperitoneally at either 10 mg/kg, 2 mg/kg or 0.1 mg/kg dose in female BALB/c mice 2 hours prior to intranasal challenge with 5 murine 50% lethal doses (mLD_50_) of A/wt shearwater/WA/1979 (H15N5) virus. At the 10 mg/kg dose, 100% survival was observed with all mAbs. Most mAbs were also completely protective at the 2 mg/kg dose, except KL-H15-3G11, for which 80% of the mice survived. MAbs KL-H15-6G8 and 07-5G01 were able to confer partial protection (40% survival) even at the 0.1 mg/kg dose, however weight loss was more pronounced in these groups compared to the ones treated with higher doses of the same mAbs (Figure 4A).

**Figure 4:**
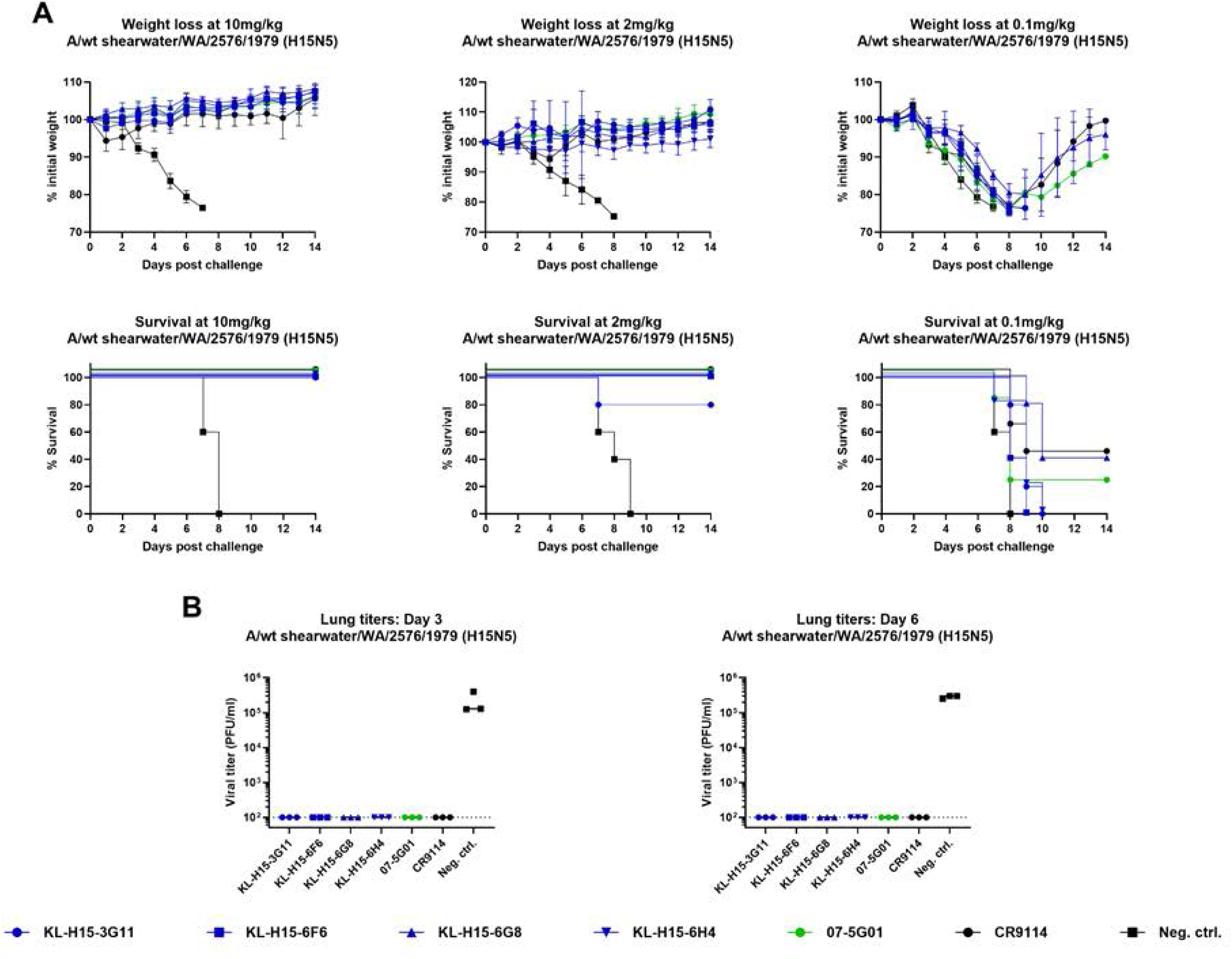
Protective efficacy of mAbs against lethal challenge *in vivo*. (A) Protection against lethal challenge in mice. Five mice per group were given mAbs intraperitoneally at a dose of 10 mg/kg, 2 mg/kg or 0.5 mg/kg and 2 hours later, they were challenged with 5 mLD_50_ of A/wt shearwater/WA/2576/1979 (H15N5) intranasally. Weight loss was monitored for 14 days post challenge and the mean weight loss for each group is shown, with error bars representing standard deviation. Survival plots for the same duration are shown. (B) Viral lung titers after infection with A/wt shearwater/WA/2576/1979 (H15N5) virus. Six mice per group were given mAbs intraperitoneally at 10 mg/kg and 2 hours later, challenged with 1 mLD_50_ of A/wt shearwater/WA/2576/1979 (H15N5) intranasally. Mice were euthanized on day 3 and day 6 post infection, the lungs were harvested and viral titer in lung homogenates was determined using plaque assay.

In addition to protection, the ability of the mAbs to inhibit viral replication in lungs of mice challenged with H15 virus was also tested. MAbs were administered at 10 mg/kg dose intraperitoneally in female BALB/c mice which were then challenged with 1 mLD_50_ of A/wt shearwater/WA/1979 (H15N5) virus. Lungs of these mice were harvested at 3 and 6 days post infection (DPI) and viral load was assessed using plaque assay. No replicating virus was detected in the lungs of mice treated with all mAbs at either time points (Figure 4B). This is consistent with the *in vivo* protection experiment and, combined with the results of neutralization assays, suggests that virus neutralization might be the primary mechanism of protection.

### Structural characterization demonstrates mAbs cluster into three distinct binding classes

Although KL-H15-6H4 and 07-5G01 were recalcitrant to negative stain electron microscopy, reconstructions of KL-H15-6G8, KL-H15-3G11, and KL-H15-6F6 revealed that all three mAbs targeted the H15 HA head (Figure S3). Sequence alignments highlighted shared and distinct features utilized by the mAbs in this study: KL-H15-6H4 and KL-H15-6F6 shared similar light-chains; KL-H15-3G11, KL-H15-6G8, and KL-H15-6H4 shared similar heavy-chains; and KL-H15-3G11 and KL-H15-6G8 shared similar light-chains (Figure S4). To characterize the relationship between complementarity-determining region (CDR) sequences and antigen engagement and to interrogate how KL-H15-6H4 was uniquely able to neutralize strains isolated after 1979, structures of A/wt shearwater/WA/2576/1979 H15 HA complexes were obtained using cryo-electron microscopy (Figure 5, Table S1).

**Figure 5.**
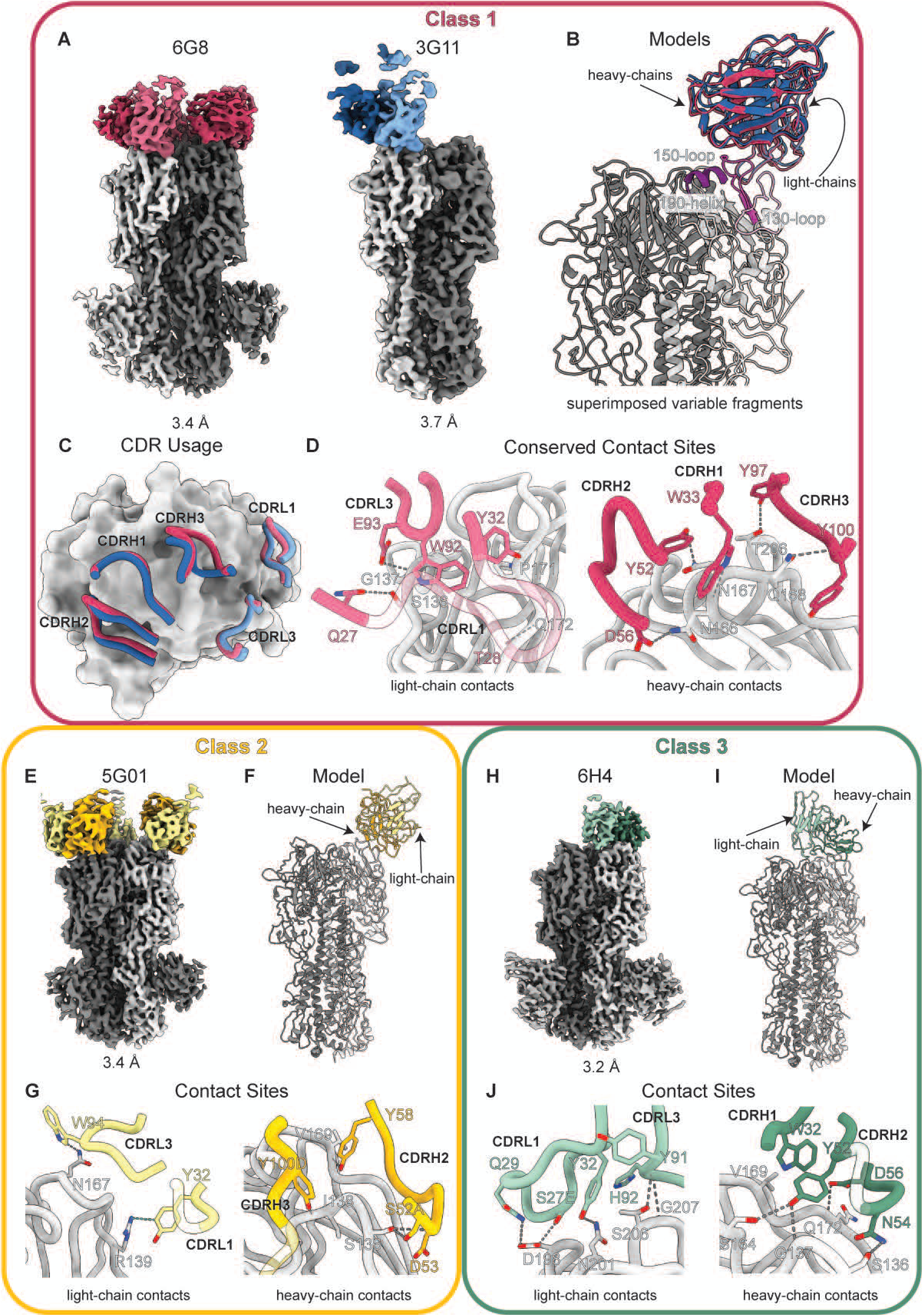
Structural characterization of Class 1, 2, and 3 epitope-paratope interactions. (A) KL-H15-6G8 and KL-H15-3G11 EM maps used for model building. (B-C) Superimposed models of KL-H15-6G8 and KL-H15-3G11 in complex with A/wt shearwater/WA/2576/1979 H15 HA highlight a shared binding mode. KL-H15-6G8 depicted in shades of red and KL-H15-3G11 in shades of blue. Heavy chains depicted in darker shades and light chains in lighter shades. The 130-loop, 150-loop, and 190-helix— antigenic regions shared with H3 HA—are depicted in shades of purple. (D) Zoomed view of light- and heavy-chain contact sites conserved across both KL-H15-6G8 and KL-H15-3G11 depicted as sticks using the KL-H15-6G8 model. Hydrogen bonds illustrated as dashed lines. All antibodies presented in Kabat numbering. (E-F) Map and model for 07-5G01 mAb binding. (G) 07-5G01 light- and heavy-chain contact sites presented as sticks with hydrogen bonds illustrated as dashed grey lines and a pi-cation interaction depicted as a light-blue dashed line. (H-I) Map and model for KL-H15-6H4 mAb binding. (J) Zoomed view of light- and heavy-chain contact sites.

As suggested by their nearly identical light- and heavy-chain CDRs, KL-H15-3G11 and KL-H15-6G8 engaged A/wt shearwater/WA/2576/1979 H15 HA in a nearly identical manner, utilizing a series of aromatic and polar uncharged residues to form a network of hydrogen bonds with H15 HA residues in or near the 130-loop, 150-loop, and 190-helix (Figure 5A-D, Figure S5A-B, Table S2). Given their similarity, KL-H15-3G11 and KL-H15-6G8 were grouped into Class 1. All heavy-chain CDRs and CDRL1 and 3 of Class 1 mAbs were involved in antigen engagement (Figure 5C). Atomic modeling of KL-H15-6G8 showed 3 hydrogen bonds formed between H15 HA and CDRL1 and CDRL3 and at least 5 formed with CDRH1, CDRH2, and CDRH3 (Fig 5D). The heavy-chain CDRs engaged residues N166, N167, Q168, and T206. Q27 in CDRL1 and E93 in CDRL3 formed hydrogen bonds with S136 and G137, respectively while T28 in CDRL1 formed a backbone-to-backbone hydrogen bond with Q172. Hydrophobic interactions between P171 on the H15 HA protomer and Y32 in CDRL1 and W92 in CDRL3 were also identified (Figure 5D).

07-5G01 engaged the H15 HA trimer in a distinct fashion and is defined as a Class 2 mAb (Figure 5E, F, Figure S5C). In 07-5G01, rather than forming a hydrophobic interaction with P171, W94 in CDRL3 formed a hydrogen bond with N167 analogous to the interaction between N167 and CDRH1 W33 in KL-H15-6G8 and KL-H15-3G11 (Figure 5G). Y32 in CDRL1 forms a pi-cation interaction with R139 rather than a van der Waals interaction with P171. In the heavy-chain, S52A in CDRH2 forms a side-chain-to-side-chain hydrogen bond with S136 while D53, Y58, and Y100D form side-chain-to-backbone hydrogen bonds with residues S163, V169, and I138, respectively.

Unlike Class 1 and 2 mAbs, KL-H15-6H4 binds towards the center of the trimer, extending the antibody footprint towards the protomer-protomer interface (Figure 5H-I, Figure S5D). In KL-H15-6H4, antigen engagement is primarily mediated by CDRL1 and CDRH2 (Figure 5J). Q29 and S27E in CDRL1 both form side-chain-to-side-chain hydrogen bonds with D198 while Y32 forms a hydrogen bond with the side-chain of N201. S206 and G207 on the HA trimer form hydrogen bonds with the backbones of residues Y91 and H92 in CDRL3, respectively. Like 07-5G01, the KL-H15-6H4 heavy-chain CDRs directly interact with V169 and S163, the former through hydrophobic interactions with W32 in CDRH1 and the latter through side-chain hydrogen bonding with N54 in CDRH2. Like Q27 in 6G8 CDRL1, Y52 in KL-H15-6H4 CDRH2 forms a hydrogen bond with G137 with an additional bond formed with S164. Lastly, D52 in CDRH2 forms a hydrogen bond with the backbone of Q172. Although CDRH3 was not predicted to make prominent contacts with antigen, it should be noted that the Y97 and Y100 sidechains in KL-H15-6H4 CDRH3 were not built into the map as there was insufficient density to do so with confidence. This may be the result of high flexibility in the region at this angle of approach. While we were unable to build an atomic model for KL-H15-6F6, docking of the predicted variable fragment (Fv) revealed that contacts with the H15 trimer were mediated by CDRH1, CDRH2, and CDRL3 (Figure S5E, Figure S6). As the KL-H15-6F6 light chain variable regions are nearly identical to that of KL-H15-6H4, this observation suggests that the contacts mediated by the KL-H15-6F6 heavy chain lead to rearrangement of the Fab angle of approach such that the CDRL1 contacts observed in KL-H15-6H4 play a less critical role in antigen engagement.

Structural analysis of antigen engagement by H15 sequence conservation and putative antigenic sites revealed that all three mAb classes engage distinct but overlapping footprints on the H15 head (Figure 6A). All three classes engage primarily with major antigenic site B. All heavy-chain contacts made by Class 1 mAbs engaged residues in site B, while E93 in CDRL3 formed a hydrogen-bond with G137, which is present in both sites A and B. 07-5G01 formed direct contacts with residues in site B, but engaged with residues S136 and I138 which straddle G137 in site A and may be able to flexibly engage with R134 in putative site E. While still primarily engaging with site B residues, the KL-H15-6H4 footprint extends towards the protomer-protomer interface and incorporates a greater number of absolutely conserved residues relative to Class 1 and 2 mAbs. Our structural models reveal that the emergence of M165R/K and N166 mutations in strains isolated after 1979 would sterically hinder and/or charge repulse engagement by Class 1 mAbs, which directly interact with residues N166 and N167, and 5G01 which interacts with residue N167 (Figure 5D and G). Associated rearrangement of neighboring residues may also contribute to loss of binding and neutralization. While KL-H15-6H4 engages spatially adjacent residues, it does not form key hydrogen bonds with N166 or N167 thereby preserving binding to strains through A/duck/Bangladesh/24704/2015 (H15N9) and neutralization through A/mallard duck/Sweden/336472/2012 (H15N5) (Figure 2A, Figure 5J).

**Figure 6.**
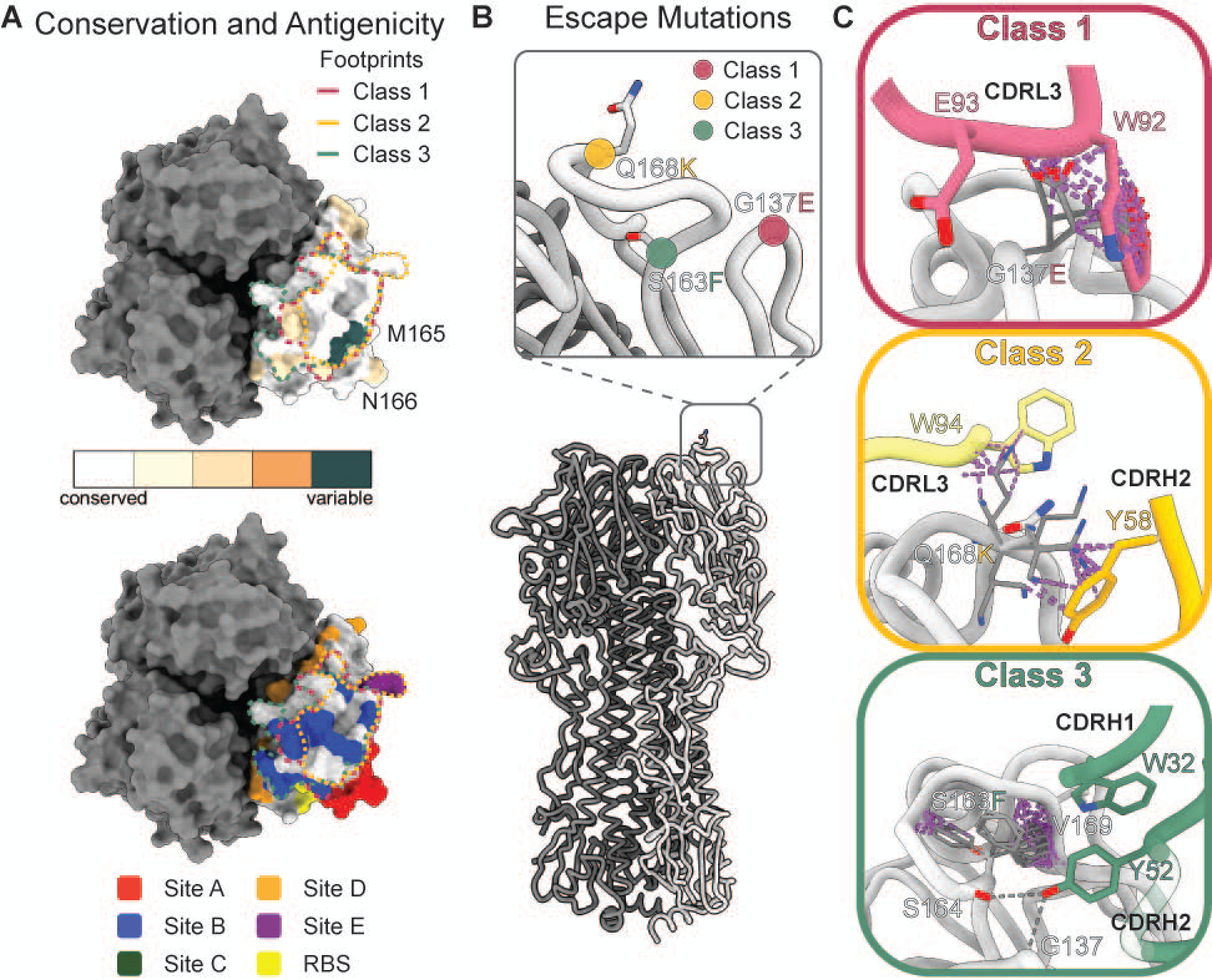
Structural analysis of mAb binding. (A) Top-view of the A/wt shearwater/WA/2576/1979 H15 trimer; bound protomer presented in light grey. Dashed lines are used to demarcate the footprints—defined as residues within 5Å of Fabs—for Class 1 in red, Class 2 in orange, and Class 3 in green. Bound protomer is colored by conservation across all 22 unique H15 sequences with degree of variability increasing from white to teal. Putative major antigenic sites are presented and color coded: Site A (red), B (blue), C (green), D (orange), and E (purple). The receptor binding site is highlighted in yellow. (B) Location of escape mutations derived from viral passage are depicted on a model of a A/wt shearwater/WA/2576/1979 H15 trimer. (C) Zoomed in views of escape mutations separated by antibody class. Mutations were modeled in ChimeraX and the top ten most prevalent rotamers are presented as sticks in grey. Steric clashes with paratope residues are depicted using purple dashed lines.

### Epitope mapping via escape mutagenesis

Given the difference in binding and neutralization activity of these mAbs against different H15 viruses tested, determining the target epitopes for these mAbs can provide insights into the evolution of H15 viruses. For the mAbs with HI activity, this was carried out in specific pathogen free (SPF) embryonated chicken eggs in a single passage in the presence of 250 µg of respective mAbs. 10^6^ plaque forming units (PFU) of A/wt shearwater/WA/1979 (H15N9) virus were incubated with either KL-H15-3G11, KL-H15-6F6, KL-H15-6G8 or 07-5G01 and injected into 10 days old embryonated chicken eggs. The allantoic fluid was harvested and screened for loss of HI activity to confirm escape. The escape mutants were plaque purified on MDCK cells, grown in embryonated chicken eggs and the HA segments were sequenced to identify the escape mutations. For KL-H15-6H4, which has been shown to be neutralizing but HI inactive, escape mutagenesis was done by sequential passaging of A/wt shearwater/WA/1979 (H15N9) virus with the mAb starting at 0.5xIC_50_ and doubling the concentration at every passage up to 128xIC_50_. The resultant viruses were screened for loss of binding, and thus escape, by immunofluorescence assay. The viruses were then plaque purified on MDCK cells, grown in embryonated chicken eggs and the HA segments were sequenced. Interestingly, the same G137E mutation was observed in the mutant viruses that escaped from KL-H15-3G11, KL-H15-6F6 and KL-H15-6G8, while an S163F mutation was identified in KL-H15-6H4 escape mutants and 07-5G01 escape mutant carried the Q168K mutation (Figure 6B, C). Both S163F and Q168K mutations are within the putative antigenic site B of the H15 HA, while the G137E mutation is within the putative antigenic site A. These findings further corroborate the neutralization potential of these mAbs. While these particular residues are conserved across different H15s tested in this study, there are several mutations in the regions flanking these sites which might explain the difference in binding and neutralization profile of these mAbs. Our structural models also clarify the molecular basis of escape mutations identified in this study: G137E (Class 1), Q168K (Class 2), and S163F (Class 3) (Figure 6B-C). G137E may disrupt hydrogen bonding between residues G137 in the trimer and E93 in CDRL1 and may introduce steric clashes with residue W92. Similar to the naturally occurring M165R/K and N166 mutations, Q168K introduces a large, positively-charged residue into a polar uncharged pocket, potentially creating steric clashes and disrupting hydrogen bonding networks mediated by W94 in CDRL3 and Y58in CDRH2. In contrast, S163F does not directly interfere with residues in the KL-H15-6H4 paratope, but analysis of steric clashes suggests that it may disrupt the epitope pocket and thereby ablate neutralization.

### Polyclonal serum responses to different H15 viruses indicate antigenic drift

The remarkable difference in the binding and neutralization activity of the mAbs tested in this study, particularly the mAbs that have HI activity against A/wt shearwater/WA/1979 (H15N9) virus hint at antigenic differences between H15 viruses isolated in different years. This can be further illuminated with the help of polyclonal responses to different H15 viruses. Groups of female BALB/c mice were infected with different H15 viruses, sera were collected 21 DPI and analyzed in an HI assay against both homologous and heterologous H15 viruses. No detectable HI response was observed against the homologous strain in sera obtained from mice infected with A/wt shearwater/WA/1979 (H15N9), hence the same mice were infected a second time, sera were collected 7 DPI and used for the HI assay. Remarkable differences were observed between serum HI reactivity against homologous and heterologous H15 viruses. Most noticeably, the sera from mice infected with A/wt shearwater/WA/1979 (H15N9) showed a significant drop in HI activity against newer H15 viruses (Table 1). This is similar to the pattern observed with the HI-active mAbs from the panel and further hints at major antigenic differences between the H15 from the 1979 reference strain to the most recent strain isolated in 2015.

**Table 1.** Serum HI cross reactivity: The values represent the last reciprocal dilution of sera at which HI activity was observed. Median values are shown in bold font, followed by the range in parenthesis.

| Strain used to infect mice | Strain used in HI assay |  |  |  |  |
| --- | --- | --- | --- | --- | --- |
|  | A/wt<br>shearwater/<br>WA/ | A/teal/Cha<br>ny/<br>7119/2008 | A/mallard/<br>Novomychali<br>vka/<br>2-23-12/2010 | A/mallard<br>duck/<br>Sweden/ | A/duck/<br>Banglade<br>sh/ |
|  | **2576/1979** |  |  | **139647/2012** | **24704/2015** |
| **A/wt shearwater/**<br>**WA/2576/1979** | **20** (10-20) | **5** (5-10) | **5** (5-10) | **5** (5-10) | **5** (5-10) |
| **A/teal/Chany/7119/2008** | **10** (5-10) | **80** (80-80) | **160** (80-160) | **40** (5-80) | **10** (10-40) |
| **A/mallard/Novomychali vka/**<br>**2-23-12/2010** | **10** (5-10) | **20** (10-80) | **320** (160-320) | **40** (10-40) | **20** (10-40) |
| **A/mallard duck/Sweden /**<br>**139647/2012** | **5** (5-5) | **5** (5-10) | **5** (5-20) | **40** (40-40) | **5** (5-10) |
| **A/duck/Bangladesh/**<br>**24704/2015** | **5** (5-5) | **10** (5-160) | **20** (10-160) | **40** (20-160) | **160** (40-320) |

## DISCUSSION

The H15 subtype of influenza A virus has rarely been detected and has so far only been found sporadically in Australia and Eurasia (El-Shesheny et al., 2018; Muzyka et al., 2012; Röhm et al., 1996; Sivay et al., 2013). Here, we investigated the antigenicity of this subtype and its different isolates using both mAbs and polyclonal serum.

MAbs generated to the prototype strain from a shearwater in 1979 which bound to putative antigenic sites A and B (as compared with H3 HA) rapidly lost binding and neutralizing activity to newer H15 virus isolates. Similarly, cross-reactivity of isolate specific sera in HI assays was relatively poor. MAbs showed strong neutralization and HI activity and protected mice potently from an H15 infection. The exception was mAb KL-H15-6H4 which had no HI activity but was the only mAb that showed breadth and bound to all tested viruses. This mAb had neutralizing activity (despite not being HI active) and also robustly protected mice from H15 virus challenge. These results suggest substantial antigenic drift over time, which is noteworthy since this subtype is only detected rarely – suggesting limited circulation, at least in sampled species.

While putative sites A and B on H15 correlate well with their H3 counterparts, sites C, D, and E do not map onto H15 so analogously (Figure 6A). Putative sites C and E are spatially discontiguous and over half of residues in putative site D are buried in relatively inaccessible regions of the protomer-protomer interface. Analysis of variation in known H15 HA sequences may offer insights into how to better characterize antigenic variation in this related but rare and distinct subtype. Notably, multiple sequences collected from Australian ducks in 1983 contain clusters of amino acid mutations in or near putative antigenic sites D and E that are not observed before or after: A228D (site D), T180N (site D proximal), L86K (site E), and S130P (site E). All except T180N constitute mutations that introduce an electric charge or remove a polar side chain and all except L86K are within or near well-characterized structural features also present in H7 and H10 HAs: loop-220, helix-190, and loop-130 (Tzarum et al., 2017; Yang et al., 2020)In addition to these mutations unique to 1983 strains, D57N (site C), K202Q (site B), and B151G (site A) are observed across virtually all strains reported after 1983—A/mallard/Novomychalivka/2-23-12/2010 being the exception as it lacks V151G. While it cannot be stated unambiguously, the location of these mutations and relatively large changes in side chain chemistry are suggestive of substantial immune pressure and antigenic drift.

Despite the variation observed in viral isolates at putative sites C, D, and E, all mAbs in this study exclusively engage sites A and B, and primarily site B. Although it has been observed that natural infection with H7N9 viruses in humans can induce antibody responses to sites A, B, the HA stem, and the trimer interface, site B is dominant in human antibody responses to circulating H3N2 strains (Gilchuk et al., 2021; Li et al., 2019; Popova et al., 2012; Wu et al., 2020). While there have been several studies on the immunogenicity of H7 and H10 in mice and ferrets, they primary utilize serological techniques that do not disambiguate the relative immunodominance of major antigenic sites (Bahl et al., 2017; Fadlallah et al., 2020; Kamal et al., 2017). Thus, it is unclear whether the site A and B dominance observed in this study is generalizable to H7 and H10 as well or if it is a feature of mouse immunization with the A/wt shearwater/WA/2576/79 (H15N5) strain.

This is a salient question, as there is a paucity of data characterizing the molecular basis of antibody responses to avian IAVs in wild or domesticated birds due to challenges inherent in working in such systems. There are multiple reasons to believe that studies of mammalian antibody responses to avian influenza viruses—and other avian pathogens for that matter—may not reflect avian responses: 1) the primary immunoglobulin in birds is IgY, which is related to but distinct from IgG; 2) in birds, IAV is primarily a gastrointestinal disease and thus the avian immune response and particularly the avian mucosal immune response likely relies on local T and B cell populations that are distinct from those involved in the mammalian response to respiratory IAV infections; and 3) as the natural reservoirs for IAVs, birds may experience repeat infections and thus studies of their antibody responses and/or potential vaccination strategies may be needed to overcome the challenge of pre-existing immunity (Causey & Edwards, 2008; Dascalu et al., 2024; Hill et al., 2016; Leslie & Clem, 1969; Meade et al., 2017; Spackman, 2009; Vandegrift et al., 2010; Zhang et al., 2017). Increasing our understanding of the immune pressures on avian IAVs will be an essential component of efforts to vaccinate critically endangered wild birds, as has already been attempted with the California condor (Katzner et al., 2025).

While influenza viruses mutate and change over time, significant changes are often driven by evolutionary pressure such as a switch in host species and adaptation to the new species or reinfection of hosts (meaning neutralizing antibodies are present that exert pressure on the virus). It is unclear which mechanisms led to these significant antigenic changes in H15 viruses. Furthermore, the rare detection suggests low prevalence, but the antigenic changes suggest that the virus is circulating in a larger reservoir. What this reservoir is, needs to be determined. However, it is clear that the virus is not widespread in *Charadriiformes*, *Procellariiformes* or *Anseriformes* due to the rare detections. Potentially, viruses detected in these birds are an indication of spill-over events from the real reservoir.

We believe our study sheds some light on this rare influenza A subtype – even though it raises several questions that need to be answered. Furthermore, the produced mAbs, recombinant proteins and re-assortant virus represent a set of reagents for the influenza virus research community to better study historic and future H15 viruses.

## RESOURCE AVAILABILITY

## Lead contact

Requests for information or reagents should be directed to Florian Krammer.

## Materials availability

mAbs and plasmids for mAb expression can be obtained from the authors upon reasonable request.

## Data and code availability

- CryoEM maps and models were deposited in the Electron Microscopy DataBank under accession IDs EMD-71419, EMD-71420, EMD-71421, EMD-71422, and EMD-71424, and in the Protein DataBank under accession IDs 9P9Q, 9P9R, 9P9S, and 9P9T (Table S1).
- The published article contains all datasets analyzed during the study.
- This paper does not report original code.
- Any additional information required to reanalyze the data reported in this paper is available from the lead contact upon request.

## ACKNOWLEDGEMENTS

This work was also partially funded by the Centers of Excellence for Influenza Research and Response (CEIRR, contract #75N93021C00014, reagent generation), the Collaborative Influenza Vaccine Innovation Centers (CIVIC, National Institute of Allergy and Infectious Diseases contract #75N93019C00051), and by institutional funds. E.P.M. was also supported by a postdoctoral fellowship from Fundación Ramón Areces. We thank Paul E. Leon for pilot study work on these mAbs. We thank Jesse Bloom for making his HA renumbering algorithm publicly available and the Bacterial and Viral Bioinformatics Resource Center (BV-BRC) for hosting it on an accessible user interface.

## AUTHOR CONTRIBUTIONS

D.B. and F.K. designed the study. D.B. and A.L. designed and performed the experiments and analyzed data. W.H., and E.P.M. performed experiments and/or generated reagents. P.C.W. provided reagents. J.H., A.B.W. and F.K. supervised the study. D.B., A.L. and F.K. drafted the manuscript. All authors reviewed, edited and approved the final version of the manuscript.

## STAR Methods

### Experimental model and study participant details

Mab 07-5G01 was generated from the sequences obtained from plasmablasts isolated from an individual who received two doses of a live-attenuated cold-adapted influenza A/Anhui/1/2013 (H7N9) vaccine followed by an inactivated virus vaccine boost 12 weeks later. The details of the sample collection are described in a previous publication: Henry Dunand, Carole J et al. “Both Neutralizing and Non-Neutralizing Human H7N9 Influenza Vaccine-Induced Monoclonal Antibodies Confer Protection.” Cell host & microbe vol. 19,6 (2016): 800-13. doi:10.1016/j.chom.2016.05.014 PMCID: PMC4901526 NIHMSID: NIHMS789757 PMID: 27281570.

### Cells, viruses and recombinant glycoproteins

Dulbecco’s modified Eagle’s medium (DMEM; Gibco) supplemented with Pen-Strep antibiotic solution (penicillin at 100 U/ml and streptomycin at 100 µg/ml; Gibco), 10 ml of 1 M 4-(2-hydroxyethyl)-1-piperazineethanesulfonic acid (HEPES; Life Technologies) and 10% fetal bovine serum (FBS; Gibco) was used to culture MDCK (ATCC# CCL34) and human embryonic kidney 293T (HEK-293T; ATCC# CRL-3216) cells in a 37°C incubator with 5% CO_2_. Expi293 medium (Gibco) was used to culture Expi293F (Gibco# A14527) cells in suspension culture shaking at 125 RPM in an incubator at 37°C and 8% CO_2_. Hybridoma clones were initially cultured in ClonaCell-HY Medium E (Stemcell Technologies) and progressively switched to culture in hybridoma-serum free medium (Hybridoma-SFM; Gibco). *Spodoptera frugiperda* (Sf9) cells were grown in TNM-FH insect medium (Gemini Bio) supplemented with 10% FBS and antibiotics (100 U/ml of penicillin, 100 µg/ml streptomycin; Gibco). BTI-TN-5B1-4 (Trichoplusia ni) cells were grown in serum-free Sf-900 II medium (Gibco) supplemented with antibiotics (100 U/ml of penicillin, 100 µg/ml streptomycin; Gibco).

HA and NA sequences from the following isolates were obtained from GenBank and were commercially synthesized (Genewiz; Azenta Life Sciences): A/wt shearwater/WA/2576/1979 (H15N9; CY006010 and CY005407), A/teal/Chany/7119/2008 (H15N4; CY098540 and CY098542), A/mallard/Novomychalivka/2-23-12/2010 (H15N7; KP087869), A/mallard duck/Sweden/139647/2012 (H15N5; MF147992 and MF148018) and A/duck/Bangladesh/24704/2015 (H15N9; KY635680 and KY635500). Recombinant H15 viruses were rescued with the HA and NA from the original isolate and the remaining six segments from A/PR/8/34 (A/PR/8/34, H1N1) as 6:2 reassortant viruses using the protocol described previously (Martínez-Sobrido & García-Sastre, 2010). The A/PR/8/34 backbone is attenuated in humans and generally considered safe (Beare et al., 1975). A/mallard/Novomychalivka/2-23-12/2010 (H15N7) virus was rescued with HA from the original isolate and the remaining seven segments from A/PR/8/34 as 7:1 reassortant virus, as the NA sequence from the original isolate is incomplete. A/wt shearwater/WA/2576/1979 (H15N5) mouse challenge virus was rescued using NA sequence from A/mallard/Sweden/86/2003 (H12N5; GenBank accession: CY060392.1). Virus stocks were grown in SPF-embryonated chicken eggs for 48 hours at 37°C followed by harvest of allantoic fluid and clarification by centrifugation at 300x*g* for 5 minutes. The virus stocks were stored at −80°C before use.

A/wt shearwater/WA/2576/1979 purified virus preparation for ELISA was made by low-speed centrifugation of allantoic fluid to remove debris (300 × g at 4°C for 5 minutes), followed by ultracentrifugation at 25,000 rpm for 2 hours at 4°C with an SW-28 rotor in a Beckman L7-65 ultracentrifuge through a 30% sucrose cushion buffered with NTE buffer (100 mM NaCl, 10 mM Tris-HCl, and 1 mM ethylenediaminetetraacetic acid EDTA balanced to pH 7.4). The supernatant containing allantoic fluid was aspirated, and the pellet was resuspended in phosphate buffered saline (PBS).

Recombinant HA proteins were expressed and purified using a mammalian expression system as previously described (Sano et al., 2021). Briefly, the ectodomains of HA proteins from A/wt shearwater/WA/2576/1979 (H15N9), A/teal/Chany/7119/2008 (H15N4), A/mallard/Novomychalivka/2-23-12/2010 (H15N7), A/mallard duck/Sweden/139647/2012 (H15N5) and A/duck/Bangladesh/24704/2015 (H15N9) were cloned into a pCXSN vector containing thrombin cleavage site, a foldon T4 trimerization domain and a hexa-histidine tag, and transiently transfected into Expi293F cells using an Expi293 transfection kit per the manufacturer’s instructions. The supernatant was collected 7 days post transfection and purified using affinity purification with nickel-nitrilotriacetic acid (Ni-NTA) agarose beads (Qiagen). Recombinant H1 from A/PR/8/34 and cH15/3 were produced using the baculovirus expression system as described in detail previously (Margine et al., 2013). The ectodomains of the HA proteins were cloned into a baculovirus shuttle vector. This shuttle vector contains a C-terminal trimerization domain as well as a hexahistidine tag. Baculoviruses were propagated in Sf9 cells and these viruses were then used to infect BTI-TN-5B1-4 cells. Proteins were purified from the supernatant as described above.

Plasmids for expression of mAb 07-5G01 were generously contributed by Dr. Patrick Wilson. The expression plasmids were transfected into Expi293F cells using an Expi293 transfection kit per the manufacturer’s instructions. The supernatant was collected 7 days post transfection and purified using CaptureSelect IgG-Fc (Multispecies) Affinity Matrix (Thermo Scientific).

### Generation and selection of hybridomas secreting H15 antibodies

Six to eight week-old female BALB/c mice were infected with a sublethal dose of A/wt shearwater/WA/2576/1979 (H15N5) virus and boosted with an inactivated preparation of the same virus. Three days after the boost, the mice were euthanized, and the spleens were harvested. The splenocytes were fused with SP2/0 myeloma cells using polyethylene glycol (PEG; Sigma-Aldrich) and the resultant fused cells were grown in selection medium (Clonacell-HY Medium D; StemCell Technologies). Ten to fourteen days later, individual colonies were picked and grown in 96-well plates. The supernatants from these wells were screened via ELISA for binding to the A/wt shearwater/WA/2576/1979 HA. The clones which showed reactivity were isotyped using a Pierce rapid antibody isotyping kit (Thermo scientific). The single clones were then further expanded using Clonacell-HY Medium E (StemCell Technologies) and later adapted to serum free medium (Hybridoma-SFM; Gibco). Once expanded to the desired volume, the supernatants were collected, and the antibodies were purified using CaptureSelect IgG-Fc (Multispecies) Affinity Matrix (Thermo Scientific).

All *in vivo* experiments were conducted in accordance with the guidelines of the Icahn School of Medicine at Mount Sinai Institutional Animal Care and Use Committee (IACUC).

## ELISA

96-well plates (Immulon-4 HBX, Thermo Scientific) were coated with 50 µl/well of recombinant protein diluted to a concentration of 2 µg/ml in PBS and stored at 4°C overnight. On the following day, the coating solution was discarded, and the plates were blocked with 200 µl/well of 3% non-fat milk prepared in PBS with 0.1% Tween-20 (PBS-T) for one hour at room temperature (RT). MAbs were diluted to a starting concentration of 30 µg/ml and serially diluted 1:3 in 1% non-fat milk prepared in PBS-T. After one hour, the blocking solution was discarded and 100 µl of the mAb dilutions were added to the plates in duplicates and incubated for 2 hours at RT. The plates were then washed 3 times with 200 µl/well of PBS-T. Next, 50 µl/well of the secondary antibodies [for mouse mAbs: Sheep anti-mouse IgG (H&L) Antibody Peroxidase Conjugated antibody (Rockland Immunochemicals), for human mAbs: Goat anti−human IgG (Fab-specific)−horseradish peroxidase (HRP) labeled antibody (Sigma)] diluted to 1:3000 in 1% non-fat milk in PBS-T was added and incubated at RT for 1 hour. The plates were washed 3 times with 200 µl/well of PBS-T. Following this, 100 µl/well of SigmaFAST OPD (o-phenylenediamine dihydrochloride; Sigma-Aldrich) was added to the plates and incubated at RT for 10 minutes. The reaction was stopped by adding 50 µl/well of 3M hydrochloric acid (HCl; Fisher Scientific) and analyzed at an optical density (OD) of 490 nm using SynergyH1 microplate reader (BioTek). For ELISA performed with purified virus, the purified virus was coated on 96-well plates at concentration of 5 µg/ml in PBS and the protocol was followed as described above.

### Immunofluorescence assay

MDCK cells were seeded in a 96-well cell culture plate (Corning) at a density of 25,000 cells/well and cultured overnight at 37°C. The viruses were diluted in 1X minimum essential medium (MEM). The cells were washed with 200 µl/well of PBS and infected with a multiplicity of infection (MOI) of 1 for each respective virus at 37°C overnight. On the following day, the cells were fixed with 100 µl/well of 3.7% paraformaldehyde for 1 hour at RT. Next, the wells were blocked with 200 µl/well of 3% non-fat milk prepared in PBS for 1 hour at RT. After removing the blocking solution, 100 µl/well of each of the mAbs diluted to 30 µg/ml in 1% non-fat milk prepared in PBS were added to the wells and incubated at RT for 1 hour. After the incubation, the primary antibodies were removed, and the cells were washed 3 times with PBS. 100 µl/well of secondary antibodies [for mouse mAbs: Goat anti-Mouse IgG (H+L) Cross-Adsorbed Secondary Antibody, Alexa Fluor 488 (Thermo Fisher Scientific), and for human mAbs: Goat anti-Human IgG (H+L) Cross-Adsorbed Secondary Antibody, Alexa Fluor 488 (Thermo Fisher Scientific)] diluted to 1 µg/ml and 4’,6-diamidino-2-phenylindole, dilactate(DAPI; Thermo Scientific) diluted to 1:500 in 1% non-fat milk in PBS were added to the wells and incubated at RT for 1 hour covered in foil. The cells were washed 3 times with PBS and imaged using EVOS M5000 fluorescent imaging microscope (Thermo Scientific).

### Western blot assay

Ten (10) ng of recombinant H15 from A/wt shearwater/WA/2576/1979 and H1 from A/PR/8/34 (negative control) prepared in PBS were mixed with equal volume of 2x Laemmli sample buffer supplemented with beta-mercaptoethanol and incubated at 95°C for 20 minutes. The proteins were then loaded and run on 4-20% gradient SDS polyacrylamide gel (Bio-Rad) and later transferred to nitrocellulose membranes using iBlot 2 Gel Transfer Device (Invitrogen). The membranes were blocked with 3% non-fat milk in PBS-T and later probed with 30 µg/ml of each mAb diluted in 1% non-fat milk in PBS-T for 1 hour at RT. Anti-6x-His antibody (Thermo Fisher Scientific) was used as a positive control. Following this, the membranes were washed 3 times with PBS-T and stained with alkaline phosphatase (AP) conjugated secondary antibody (Sigma) for 1 hour at RT. The blots were developed with AP conjugate substrate kit (Bio-Rad) according to the manufacturer’s protocol.

### Microneutralization assay

MDCK cells were seeded in 96-well cell culture plates at a density of 25,000 cells/well and cultured overnight at 37°C. The viruses were diluted to 100x 50% tissue culture infectious dose (TCID_50_)/50 µl and three-fold serial dilutions of mAbs starting at 30 µg/ml in duplicates were prepared in 1X MEM supplemented with 1 µg/ml tosyl phenylalanyl chloromethyl ketone (TPCK)-treated trypsin (Sigma). Antibody dilutions and viruses were incubated together for 1 hour at RT on a shaking platform. The cells were washed with PBS and the antibody-virus mix was added and incubated at 37°C for 1 hour. Another set of mAb dilutions were prepared in 1X MEM supplemented with 1 µg/ml TPCK-treated trypsin. At the end of the incubation, the cells were washed with PBS and mAb dilutions prepared in 1X MEM with TPCK-treated trypsin were added. The cells were then incubated at 37°C for 48 hours. At the end of the incubation, hemagglutination assays were performed using the supernatant from infected cells and 0.5% chicken red blood cells (RBCs; Lampire Biological Laboratories) diluted in PBS to test for the presence of virus in each well.

### Hemagglutination inhibition assay (HI)

MAbs were serially diluted 1:2 at a starting concentration of 30 µg/ml in PBS in v-bottom 96-well plates (Thermo Scientific). A hemagglutination assay was performed to determine the hemagglutination titer (HAU) for the viruses. Twenty five µl of the mAb dilutions were incubated with 25 µl of viruses diluted to 8 HAU for 1 hour at RT. Following this, 50 µl of 0.5% chicken RBCs diluted in PBS were added to the wells and incubated at 4°C for 1 hour and analyzed.

For the HI assay performed with mouse serum, sera were treated with receptor destroying enzyme (RDE; Denka-Seiken) to remove non-specific inhibitors of hemagglutination. Starting with an initial 1:10 dilution, the sera were serially diluted 1:2 in PBS in V-bottom 96-well plates. Twenty five µl of serum dilutions were incubated with 25 µl of viruses diluted to 8 HAU for 1 hour at RT. Following this, 50 µl of 0.5% chicken RBCs diluted in PBS was added to the wells and incubated at 4°C for 1 hour and analyzed.

## PRNA

MDCK cells were seeded in 12-well cell culture plates (Corning) at a density of 250,000 cells/well and cultured overnight at 37°C. 1:5 serial dilutions of mAbs starting at 100 µg/ml were prepared in 1X MEM and incubated with 1600 PFU/ml of A/wt shearwater/WA/2576/1979 (H15N9) virus for 1 hour at RT with shaking. The virus-antibody mixture was then transferred on MDCK cells in duplicates and incubated at 37°C for 1 hour with intermittent shaking every 10 minutes to prevent drying out of cells. Following the incubation, agar overlays containing corresponding mAb dilutions were applied and the cells were incubated at 37°C for 48 hours. At the end of the 48 hours, the cells were fixed using 3.7% paraformaldehyde for 1 hour at RT. The plates were blocked with 3% non-fat milk in PBS. Plaques were stained using anti-H15 mouse polyclonal serum diluted to 1:1000 in 1% non-fat milk in PBS for 1 hour at RT. Following PBS washes, secondary antibody (sheep anti-mouse IgG (H&L)-Peroxidase Conjugated antibody; Rockland Immunochemicals) diluted to 1:3000 in 1% non-fat milk in PBS was added to the cells and incubated for 1 hour at RT. Cells were then washed again with PBS and 300 µl/well of TruBlue Peroxidase substrate (Seracare) was added to visualize and count the plaques. Percent inhibition of number of plaques was calculated in comparison to a no-antibody control, and the IC50, defined as the concentration of the mAb at which there was a 50% reduction in number of plaques, was calculated on GraphPad Prism 10 using a nonlinear regression model.

### ADCC reporter assay

*In vitro* ADCC assay reporter kit (Promega) was used to determine the ADCC activity of the mAbs. MDCK cells were seeded in 96-well white cell culture plates (Nunc) at a density of 25,000 cells/well and cultured overnight at 37°C. The cells were infected with A/wt shearwater/WA/2576/1979 (H15N9) virus at an MOI of 1 in 1x MEM and incubated overnight at 37°C. 1:3 serial dilutions of mAbs at a starting concentration of 30 µg/ml were prepared in Roswell Park Memorial Institute (RPMI)-1640 medium (Gibco). Following the overnight incubation, the virus inoculum was removed from the cells. Twenty five µl of RPMI-1640 medium supplemented with low IgG FBS (Gibco), 25 µl of antibody dilutions and 25 µl of effector cells (75,000 cells/well) were added to each well and the plates were incubated for 6 hours at 37°C. At the end of incubation, 75 µl of luciferase substrate was added to each well and the plates were incubated for 10 minutes at RT covered in foil. Following this, luminescence was captured as readout using the SynergyH1 microplate reader.

## ELLA

ELLAs were performed to assess the neuraminidase activity of the viruses. 96-well plates (Immulon-4 HBX, Thermo Scientific) were coated with 100 µl/well of fetuin (Sigma) at a concentration of 25 µg/ml in 1X coating buffer (Seracare) and stored at 4°C overnight. The plates were incubated at 37°C with 2-fold serial dilutions of the viruses starting at a 1:10 initial dilution overnight. On the following day, the plates were washed 6 times with PBS-T and were incubated with 100 µl/well of 5 µg/ml peanut agglutinin conjugated to horseradish peroxidase (PNA-HRP; Sigma) for 2 hours at RT. Following this, the plates were washed 3 times with PBS-T. Hundred µl/well of SigmaFAST OPD was added to the plates and incubated at room temperature for 10 minutes. The reaction was stopped by adding 50 µl/well of 3M HCl and analyzed at OD 490nm using SynergyH1 microplate reader. The data was analyzed using GraphPad Prism 10 and 50% effective concentration (EC_50_) was determined for each virus.

To measure neuraminidase inhibition activity of the mAbs, the mAbs were serially diluted 1:3 at a starting concentration of 30 µg/ml and added to fetuin coated plates. An equal volume of viruses diluted to 2xEC_50_ were added to the mAb dilutions and incubated at 37°C for 16-18 hours. The remainder of the assay was performed as described above. Broadly reactive neuraminidase antibody 1G01 (Stadlbauer et al., 2019) or oseltamivir phosphate (U.S. Pharmacopeia) was used as a positive control.

### *In vivo* mouse challenge studies

Six to eight weeks old female BALB/c mice (n=5 per group) were injected with either 10 mg/kg, 2 mg/kg or 0.1 mg/kg of the mAbs intraperitoneally (IP). Two hours after administration of mAbs, the mice were infected with 5 mLD50 of A/wt shearwater/WA/2576/1979 (H15N5) intranasally (IN) under anesthesia and weights were monitored daily for 14 days. Loss of 25% of body weight led to humane euthanasia and the mouse was scored dead. The weight loss and survival curves were plotted using GraphPad Prism 10.

To assess the effect of the mAbs on viral replication in the lungs of infected mice, 6-8 weeks old female BALB/c mice (n=6 per group) were injected with 10 mg/kg of the mAbs IP. Two hours after administration of mAbs, the mice were infected with one mLD50 of A/wt shearwater/WA/2576/1979 (H15N5) IN under anesthesia. 3 Three mice were euthanized 3 DPI and their lungs were harvested. The remaining 3 mice in each group were euthanized 6 DPI and their lungs were harvested. The lungs were homogenized using BeadBlaster (Benchmark Scientific) and the supernatants were analyzed for viral load using standard plaque assay.

To assess the serum HI responses elicited response to infection, 6-8 weeks old female BALB/c mice (n=5 per group) were infected with 10^5^ PFU (in 50 µl of PBS) of each of the five H15 virus strains IN under anesthesia. Serum was collected at 21 DPI and tested in an HAI assay against matched and heterologous H15 virus strains. Mice in the group infected with A/wt shearwater/WA/2576/1979 (H15N9) virus were re-infected at 28 days post initial infection and sera were collected 7 days post second infection for analysis in HI assay.

### Generation of escape mutant variants

For mAbs with HI activity, 10^6^ PFU of A/wt shearwater/WA/2576/1979 (H15N9) virus were mixed with 250 µg of mAb and incubated at RT for 1 hour. Following the incubation, the virus-mAb mixture was injected into 10-days old SPF embryonated chicken eggs (Charles River Laboratories) and incubated at 37°C for 48 hours. The allantoic fluid was harvested from the eggs at the end of incubation period and tested in HI assays for loss of HI activity. The variants against which the corresponding mAbs showed complete lack of HI activity were plaque purified on MDCK cells and grown in 10-days old SPF embryonated chicken eggs. Alternatively, for neutralizing mAb without HI activity, A/wt shearwater/WA/2576/1979 virus was incubated with 0.25xIC_50_ of mAb diluted in 1x MEM supplemented with 1 µg/ml TPCK-treated trypsin at RT for 1 hour and used to infect MDCK cells seeded in 12-well plates at 33°C for 72 hours. At the end of the incubation, the supernatant was collected and a 1:10 dilution was incubated with 0.5xIC_50_ of mAb for 1 hour at RT and used to infect MDCK cells. The virus was serially passaged until 128xIC_50_ of mAb was reached. The supernatant from the final passage was used in an immunofluorescence assay to confirm loss of binding. The variants were then plaque purified on MDCK cells and grown in 10-days old SPF embryonated chicken eggs. RNA was extracted from the allantoic fluid using E.Z.N.A. viral RNA extraction kit (Omega Biotek) and cDNA was generated using the SuperScript III First Strand Synthesis System (Invitrogen). HA segments were sequenced using Sanger sequencing to identify the escape mutations. Wild type virus passaged in the presence of irrelevant mAb KL-H6-8H9 was used as negative control to distinguish the escape mutations from other mutations that may arise from adaptation to growth in cell culture or eggs.

### Sequencing variable regions of mAbs

Variable regions of the murine mAbs were sequenced using Switching Mechanism At 5’ End of RNA Template (SMARTer 5’) Rapid Amplification of cDNA Ends (RACE) kit (Takara Bio). Total RNA from hybridoma cells was extracted using TRIzol reagent (Invitrogen). cDNA was generated by reverse transcription using isotype specific constant gene 3’ primers, the SMARTer II A Oligonucleotide and SMARTScribe reverse transcriptase. The cDNA was then amplified using SeqAmp DNA polymerase (Takara Bio) using a universal 5’ primer mix and a nested 3’ primer. The purified PCR products were Sanger sequenced using the constant gene 3’ primers and the obtained sequences were entered into the international ImMunoGeneTics information system/V-QUERy and Standardization (IMGT/V-QUEST) tool (https://www.imgt.org/IMGTindex/V-QUEST.php) to verify the complete variable region sequence. For the cases with insufficient sequencing quality, the PCR products were cloned into pRACE vector (Takara Bio) using In-Fusion Snap Assembly mix (Takara Bio). The plasmids were amplified in Stellar competent cells (Takara Bio) and miniprepped using QIAPrep Spin Miniprep Kit (Qiagen) prior to Sanger sequencing and analysis using IMGT/V-QUEST. Sequences for 07-5G01 variable regions were obtained from GenBank (heavy chain: KU987561.1, light chain: KU987562.1).

### IgG digestion

Papain was activated by 15 min incubation at 37°C in a freshly prepared solution of 100 mM Tris, 2 mM EDTA, 10 mM L-cysteine, and 1 mg/ml papain. Purified monoclonal IgG were digested at 37°C for 4-5 h with activated papain at a ratio of 40 µl activated papain solution per mg of IgG to generate Fab. Iodoacetamide was added to a final concentration of 0.03 M to halt digestion prior to concentration in a 10 kDa Amicon concentrator. Fab was separated from Fc and undigested IgG by incubation with IgG Fc Capture Select Multispecies (Thermo Fisher).

### Immune complexing and negative stain EM

Fabs were incubated with A/wt shearwater/Western Australia/2576/1979 H15 HA at a 3:1 molar ratio for 1 hour at RT. Carbon-coated 400 mesh copper grids (Electron Microscopy Services) were glow-discharged (PELCO easiGlow, Ted Pella Inc.) and 3 µl of immune complexes were applied at a concentration of ∼10-20 μg/ml. Excess sample was blotted with Whatman filter paper and 3 μl of 2% w/v uranyl formate were applied to stain complexes for 60 seconds (s) twice. Micrographs were collected using Leginon and were imaged on a Tecnai F20 (FEI) with a TemCam F416 CMOS (Tietz Video and Image Processing Systems GmbH) at 120 kV, 63,000 × magnification, and 1.67 Å/pixel (Suloway et al., 2005). For each complex, 15k to 100k particles were picked and stacked using Appion and subsequently processed using Relion 3.0 or Relion 4.0b1 (Kimanius et al., 2021; Lander et al., 2009; Scheres, 2012; Suloway et al., 2005; Zivanov et al., 2018). UCSF ChimeraX was used to generate composite 3D reconstructions (Pettersen et al., 2021).

### CryoEM grid preparation

For cryo-electron microcopy, A/wt shearwater/Western Australia/2576/1979 H15 was functionalized with sulfo-NHS-SS-diazirine (Sulfo-SDAD) (Sigma Aldrich) using N-hydroxysuccinimide (NHS)-ester chemistry as described previously and per commercial protocols (Torrents de la Peña et al., 2023). Briefly, A/wt shearwater/Western Australia/2576/1979 H15 in PBS was diluted to 1 mg/ml (∼5.5 μM) and incubated with 40-fold molar excess of freshly prepared Sulfo-SDAD solution in PBS for 30 minutes at RT. To stop the NHS-ester reaction, 1M Tris-HCl, pH 8.0 was added to the solution at a final concentration of 75 mM and the sample was incubated for 5 minutes at RT. Excess crosslinker was removed using a 30 kDa Amicon concentrator.

To crosslink immune complexes, Fab were incubated with functionalized A/wt shearwater/Western Australia/2576/1979 H15 for 30-60 minutes to facilitate binding and the Sulfo-SDAD diazirine chemistry was subsequently activated by exposing the sample to a ultraviolet (UV) dose of 1461.0 mJ/cm^2^ using a UVP Crosslinker CL-3000 (Analytik Jena). To minimize trimer degradation prior to freezing, A/wt shearwater/Western Australia/2576/1979 H15 was stabilized by activating Sulfo-SDAD diazirine chemistry prior to incubation with HL-H15-3G11 Fab. To improve tumbling, 07-5G01, KL-H15-6H4, and KL-H15-6G8 were each separately co-incubated with CR9114 (Dreyfus et al., 2012) Fab and A/wt shearwater/Western Australia/2576/1979 H15 prior to cross-linking at a molar ratio of 3:3:1. KL-H15-6F6 Fab was co-incubated with CR9114 (Dreyfus et al., 2012) Fab, and A/wt shearwater/Western Australia/2576/1979 H15 at a molar ratio of 3:3:3:1 prior to cross-linking.

Quantifoil R1.2/1.3 Cu, 300-mesh grids (Quantifoil Micro Tools GmbH) were plasma treated with a PELCO easiGlow (Ted Pella Inc.) for 25 s at 15 mA immediately before use. Grids were prepared using a Vitrobot Mark IV (Thermo Fisher Scientific) set to RT, blot force 1, 100% humidity, wait time 5 s, and blot times between 3-6 s. Immune complexes with final concentrations 0.65-0.75 mg/ml were mixed with n-octyl-beta-D-glucoside (OBG) (Anatrace) to a final detergent concentration of 0.1% immediately before grid application.

### Cryo-EM data collection

The 07-5G01, KL-H15-6H4, and KL-H15-6G8 immune complexes were screened and collected on a 300 kV Titan Krios (Thermo Fisher) equipped with a Bioquantum K3 direct electron detector (Gatan). Automated data collection was conducted using Leginon at a nominal magnification of 105,000×, a pixel size of 0.833 Å, an approximate exposure dose of 50 e- /Å2 442, and a nominal defocus range of −0.8 to −1.5 μm (Suloway et al., 2005). Data sets contained 2500-4500 micrographs. For each data collection, roughly half of the micrographs were collected at 45° tilt to overcome orientation bias.

The KL-H15-3G11 and KL-H15-6F6 immune complexes were screened and collected on a 200 kV Glacios (Thermo Fisher) equipped with a Falcon IV direct electron detector (Thermo Fisher). For KL-H15-3G11, EPU (Thermo Fisher) was used for automated data collection at 190,000× nominal magnification and a pixel size of 0.725 Å, with an approximate exposure dose of 45 e- /Å2 442 and a nominal defocus range of −0.7 to −1.4 μm. Data for the 6F6 immune complex was collected using similar parameters, but on a Glacios with a pixel size of 0.718. Data sets contained 3000-6500 micrographs each.

### Cryo-EM data processing

For 07-5G01, KL-H15-6H4, and KL-H15-6G8, Appion was used for image preprocessing and UCSF MotionCor2 was used for alignment and dose weighting (Lander et al., 2009). For KL-H15-3G11 and KL-H15-6F6, CryoSPARC Live was used for image preprocessing including alignment, motion correction, and CTF estimation (Punjani et al., 2017).

Initial processing for all immune complexes was done using CryoSPARC v3.0. A blob picker was used to pick initial particles for extraction. Following multiple rounds of 2D classification, clean particle stacks were used to generate templates for template picking. Following template picking and extraction, *Ab-Initio Reconstruction* was used to generate a starting map and to continue cleaning particle stacks. This was followed either by *Heterogeneous Refinement* if necessary to improve initial resolution and alignment or a *Homogenous Refinement* if the initial *Ab-Initio Reconstruction* was already homogenous and well-aligned. For KL-H15-6H4 and KL-H15-3G11, subsequent rounds of *Homogenous Refinement* and *Non-uniform Refinement* were sufficient to generate maps of sufficient quality for model building. For 07-5G01, the *Topaz* neural network pipeline was used to improve particle picking (Bepler et al., 2019). Briefly, intensive *2D Classification* and *Rebalance 2D* were used to generate the highest quality templates for *Topaz Training*. *Topaz Extract* was used to pick high-quality particles which were subsequently extracted for use in an *Ab-Initio Reconstruction* followed by iterative rounds of *Heterogeneous*, *Homogenous*, and finally *Non-uniform Refinements*. For KL-H15-6F6 and KL-H15-6G8, particles from a *Non-uniform Refinement* job resulting from a similar processing pathway as KL-H15-6H4 and KL-H15-3G11 were transferred to Relion 4.01b (Kimanius et al., 2021). These particles were further processed using Relion 4.01b *3D Autorefine* and *CTF Refinement* before undergoing C3 symmetry expansion and focused *3D Classification* to isolate bound H15 HA protomers. Bound particles were used for a final round of *3D Autorefine* and *Post-processing* to generate final maps for model building. Final maps were deposited to the EMDB with accession codes found in Table S1.

### Atomic model building

A crystal structure of A/wt shearwater/Western Australia/2576/1979 H15 HA (PDB: 5TG8) was used as a starting model for the trimer and ABodyBuilder 2.0 was used to generate starting models of 07-5G01, KL-H15-6H4, KL-H15-3G11, KL-H15-6G8, and KL-H15-6F6 (Abanades et al., 2023; Tzarum et al., 2017). Initial models of trimer and Fab were rigid body docked into the corresponding EM maps in ChimeraX and manually refined using Coot (Emsley & Cowtan, 2004). Iterations of Phenix real-space refinement and manual refinement in Coot were used to improve model statistics (Liebschner et al., 2019). Map-to-model fit was validated using Molprobity and EMRinger in Phenix (Barad et al., 2015; Chen et al., 2010). Final models were deposited to the PDB with accession codes found in Table S1. Figures containing maps and/or EM models were generated using ChimeraX. All H15 models are in H15 numbering, and all Fabs are deposited using the Kabat numbering scheme.

### Phylogenetic analysis

Amino acid sequences of the respective HA proteins were obtained from GenBank. Alignment of the sequences was performed using Clustal Omega and the phylogenetic tree was built using the neighbor-joining tree method. The tree was finally visualized and labeled using Figtree v1.4.4.4.

### Quantification and statistical analysis

Statistical analysis was performed using GraphPad Prism 10 using non-linear regression for Figure 2 and 3, and survival analysis was visualized using Kaplan-Meier curves for Figure 4.

## SUPPLEMENTAL FIGURE TITLES AND LEGENDS

**Figure S1:**
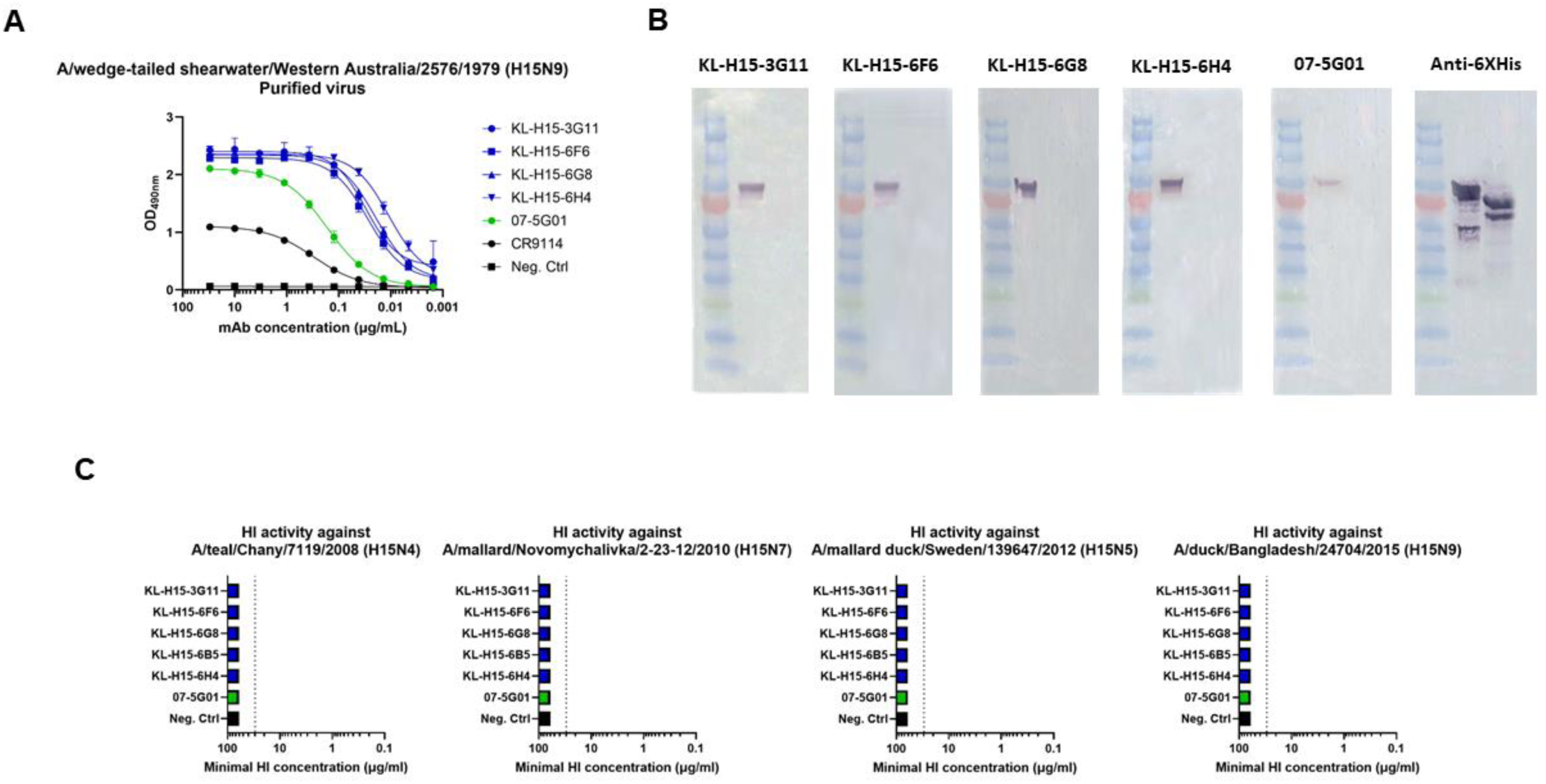
Binding and HI activity of mAbs. (A) Binding of mAbs to purified A/wt shearwater/WA/2576/1979 virus assessed by ELISA. The broadly cross-reactive HA antibody CR9114 (Dreyfus et al., 2012) was used as a positive control and anti-H6 antibody KL-H6-8H9 was used as negative control. (B) Binding to A/wt shearwater/WA/2576/1979 recombinant H15 in Western blot following a reducing denaturing SDS-PAGE. A/PR8/34 H1 was used to test for nonspecific binding, and an anti-hexahistidine antibody was used as a positive control. (C) HI activity of mAbs measured against panel of H15 viruses from 2008 to 2015. Minimal concentration at which HI activity was observed is shown here. Anti-H6 antibody KL-H6-8H9 was used as negative control.

**Figure S2:**
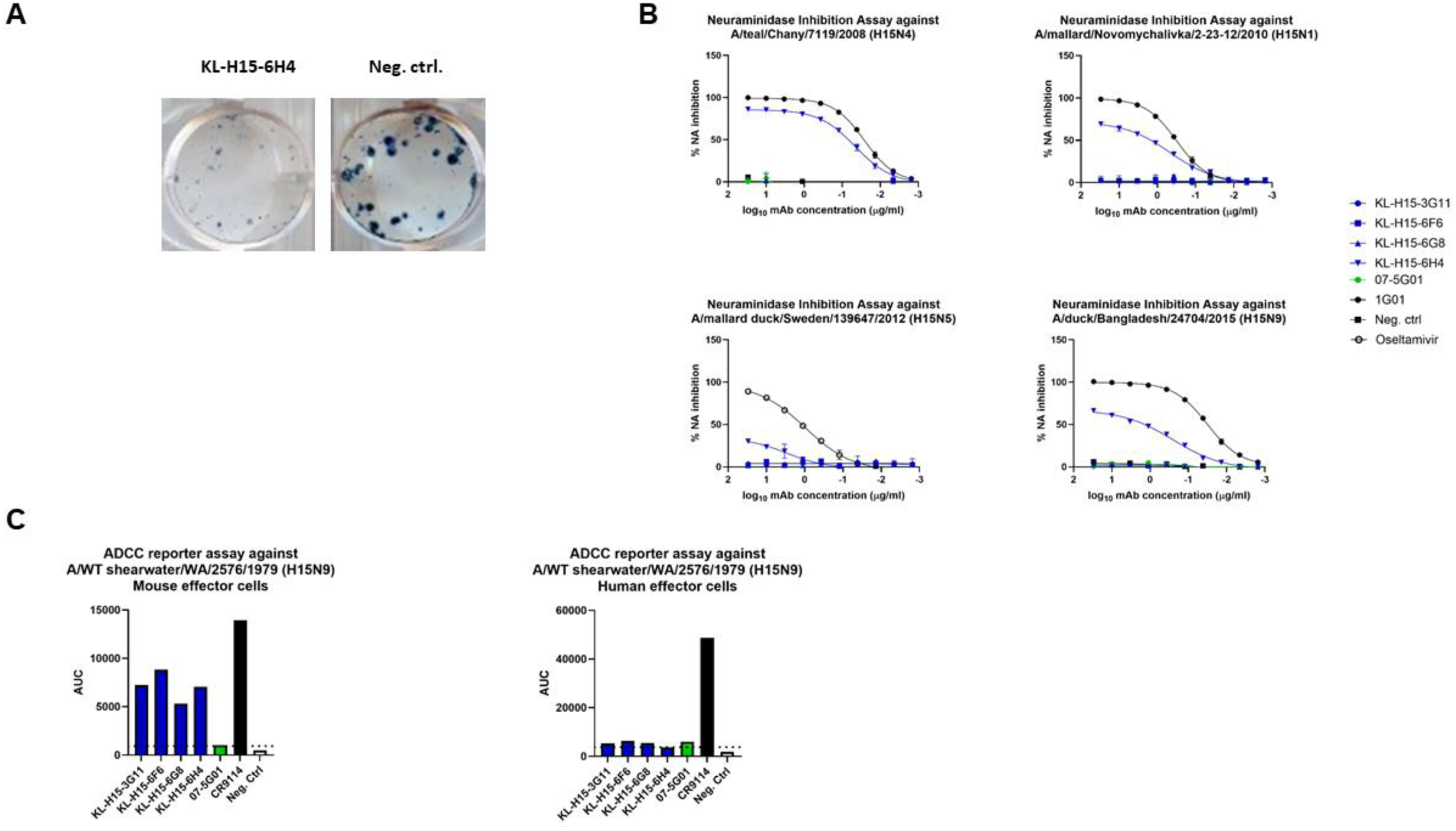
Plaque size reduction, NAI and ADCC activity of mAbs. (A) Reduction in plaque size observed in PRNA with A/wt shearwater/WA/2576/1979. Shown here are the plaques observed in the well treated with the lowest measured concentration of mAb KL-H15-6H4 as a representative of the treatment with neutralizing mAbs. The plaques observed in wells treated with anti-H6 antibody KL-H6-8H9 used as negative control are shown for comparison. (B) NAI activity of mAbs measured against panel of H15 viruses from 2008 to 2015 measured by ELLA. Broadly cross-reactive influenza virus neuraminidase antibody 1G01 (Stadlbauer et al., 2019) was used as a positive control, while anti-H6 antibody KL-H6-8H9 was used as negative control. Neuraminidase inhibitor oseltamivir was used as a positive control for NAI against A/mallard duck/Sweden/139647/2012 virus. (C) ADCC activity of mAbs was measured using antigen-specific ADCC reporter assay (Promega) using both mouse and human ADCC effector cells and the area under the curve (AUC) values are shown here. The broadly cross-reactive HA antibody CR9114 (Dreyfus et al., 2012) was used as a positive control and anti-H6 antibody KL-H6-8H9 was used as negative control.

**Figure S3:**
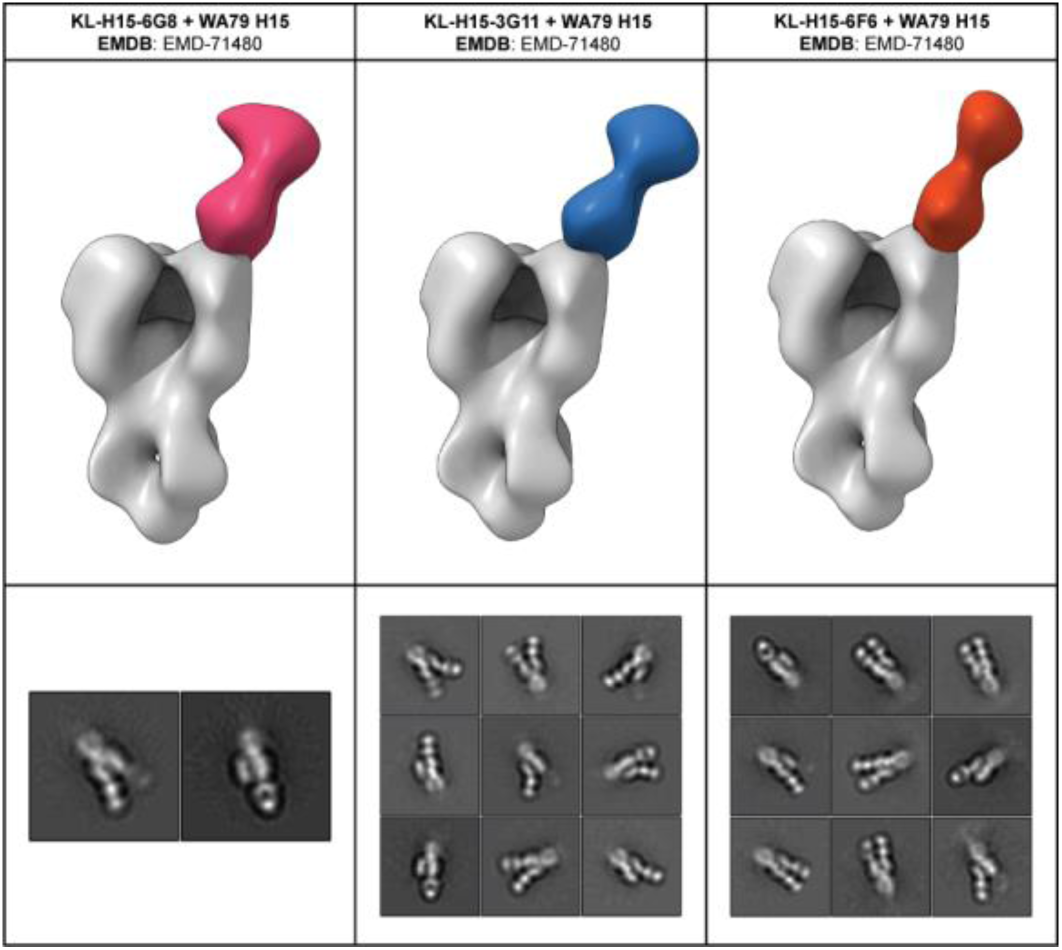
Negative stain EM maps of KL-H15-6G8, −3G11, and –6F6 Fabs complexed with A/wt shearwater/WA/2576/1979 H15 HA. Segmented Fabs from Relion 3.0 reconstructions were colored by mAb and displayed on a negative stain map of A/Hong Kong/4801/2014 H3 as a representative Group 2 HA. Representative 2D class averages of particles used to generate reconstructions are displayed below the corresponding map.

**Figure S4:**
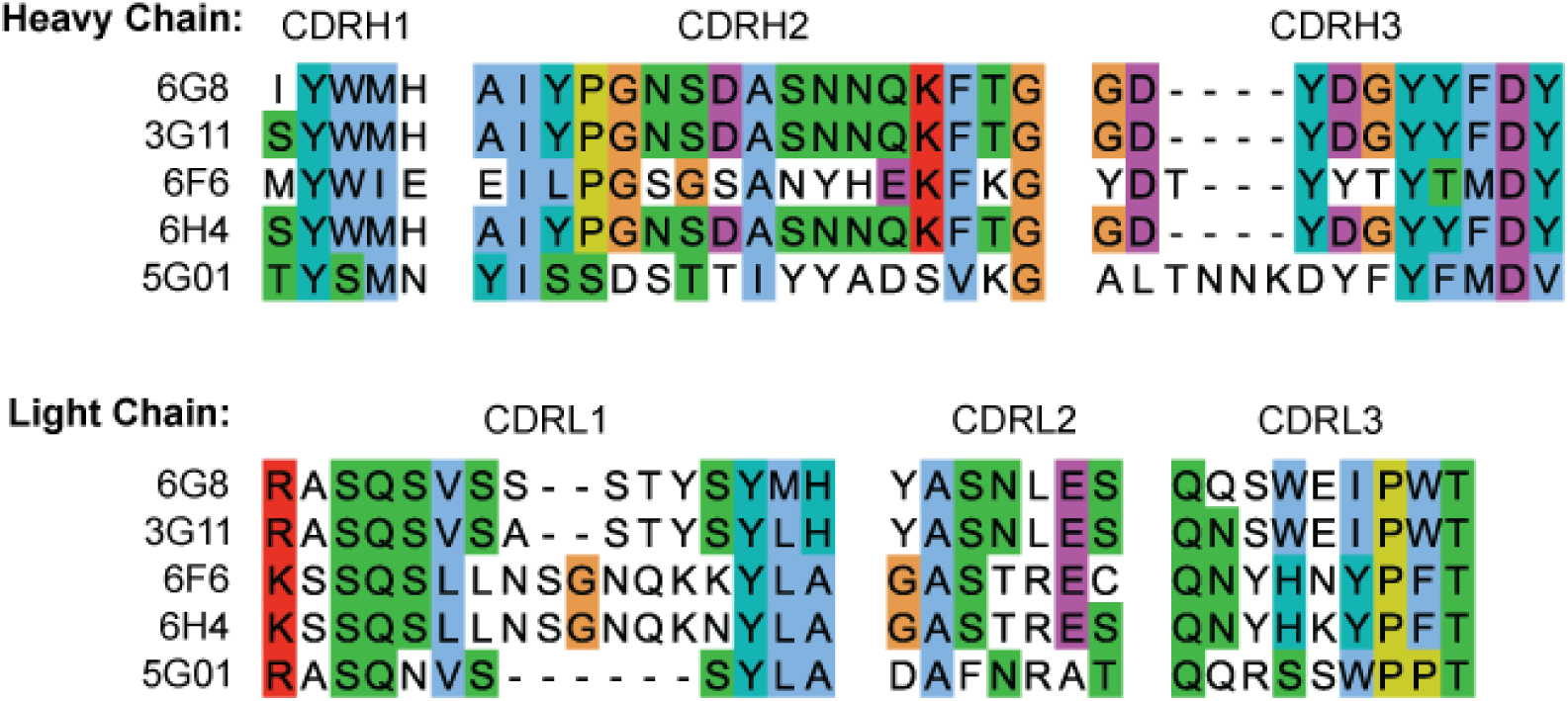
ClustalW multiple sequence alignment of heavy-chain and light-chain complementarity determining regions (CDRs) for all five mAbs reveal conserved regions within and across mAb classes. Residues are colored by conservation of side chain chemistry using ClustalW conventions.

**Figure S5:**
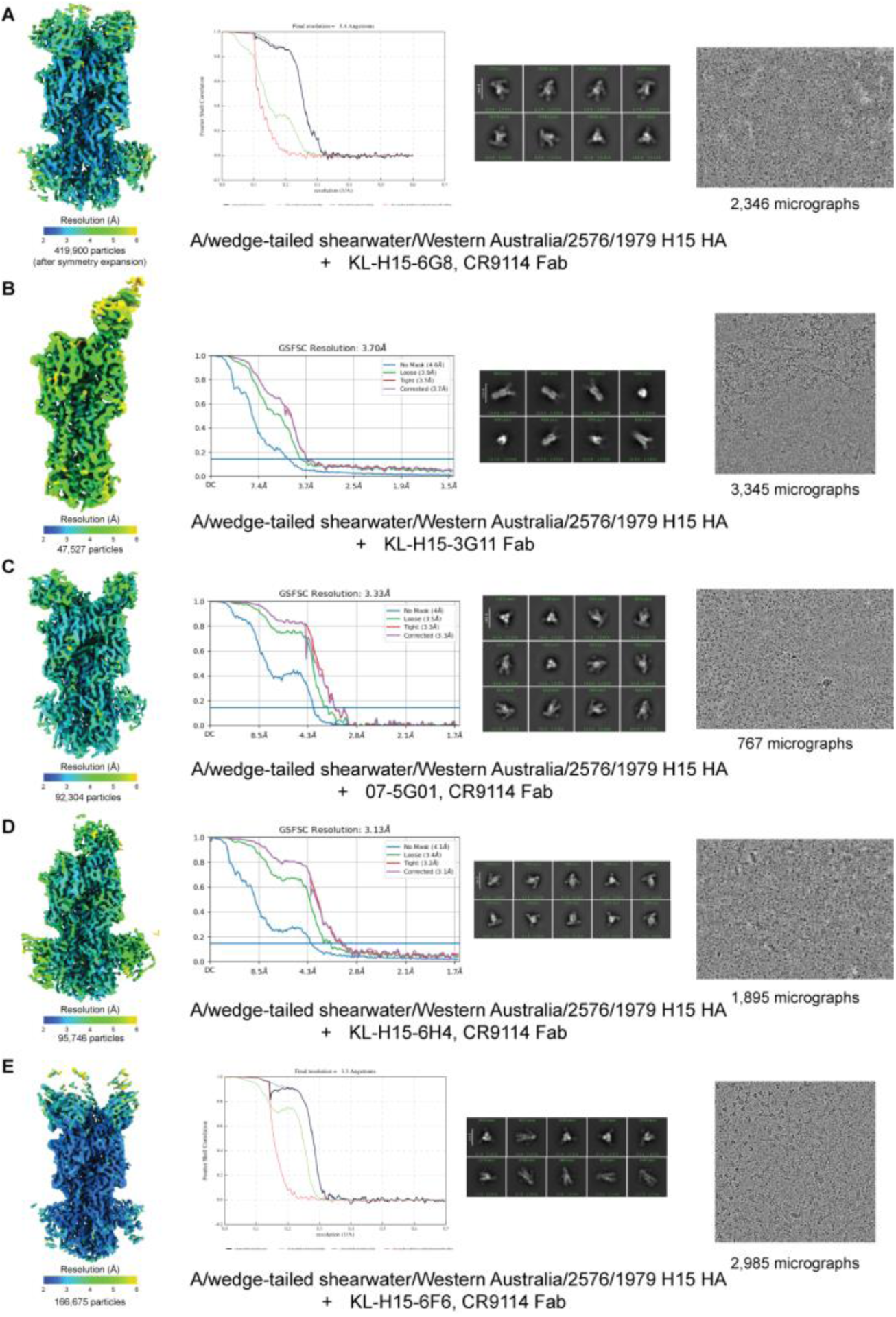
CryoEM maps, Fourier shell correlation (FSC) plots, representative 2D classes, and representative micrographs for mAb complexes. All five maps presented sequentially in rows A-E, colored by local resolution. FSC maps generated in Relion 4.01b (A and E) or CryoSPARC v4 (B-D) with resolutions reported to coincide with gold-standard FSC cutoff 0.143. All 2D classes were generated in cryoSPARC v4.

**Figure S6:**
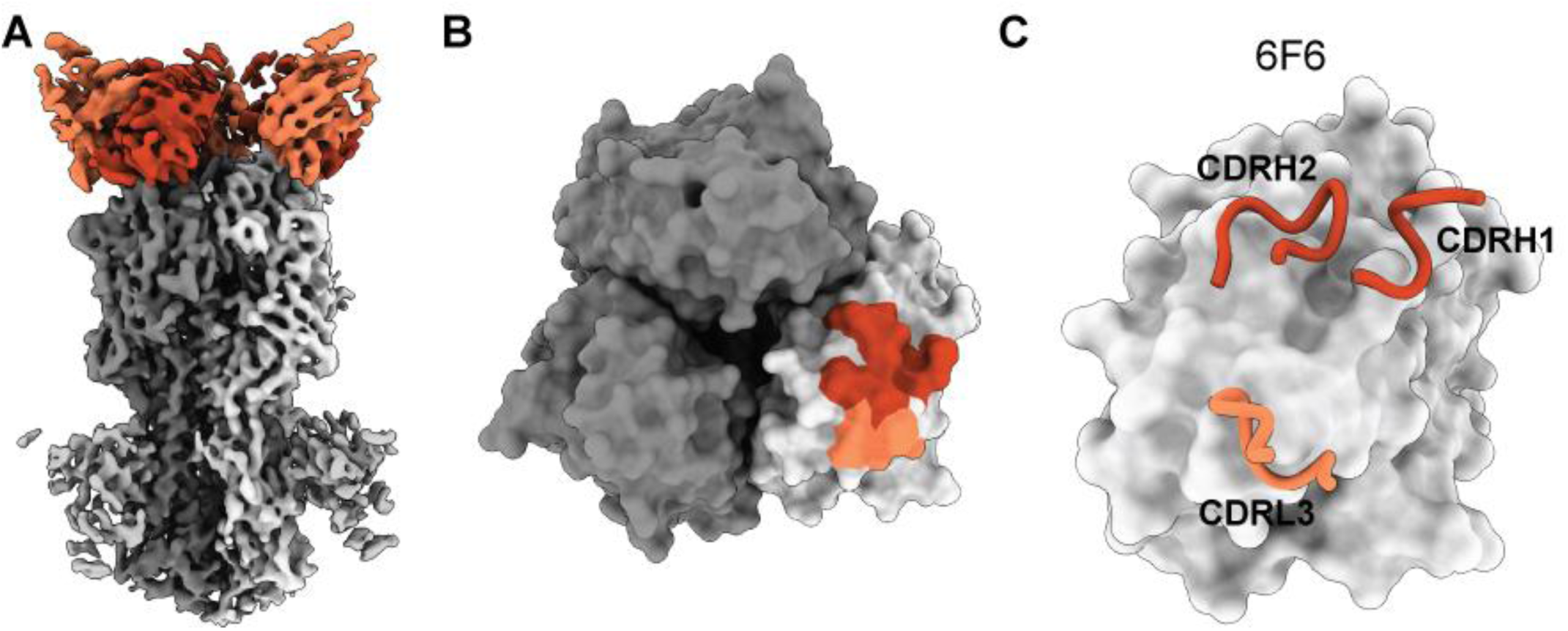
Map, footprint, and complementarity determining region usage for KL-H15-6F6 mAb. (A) Final KL-H15-6F6 EM map colored by proximity to A/wt shearwater/WA/2576/1979 H15 HA and KL-H15-6F6 Fv docked into density using ChimeraX. Light-chain depicted in light orange and heavy-chain depicted in burnt orange. (B) KL-H15-6F6 footprint on surface depiction of A/wt shearwater/WA/2576/1979 H15 HA. (C) CDRs involved in antigen engagement as demonstrated by EM map and GetContacts program (https://getcontacts.github.io/index.html, Table S2).

**Table S1.**
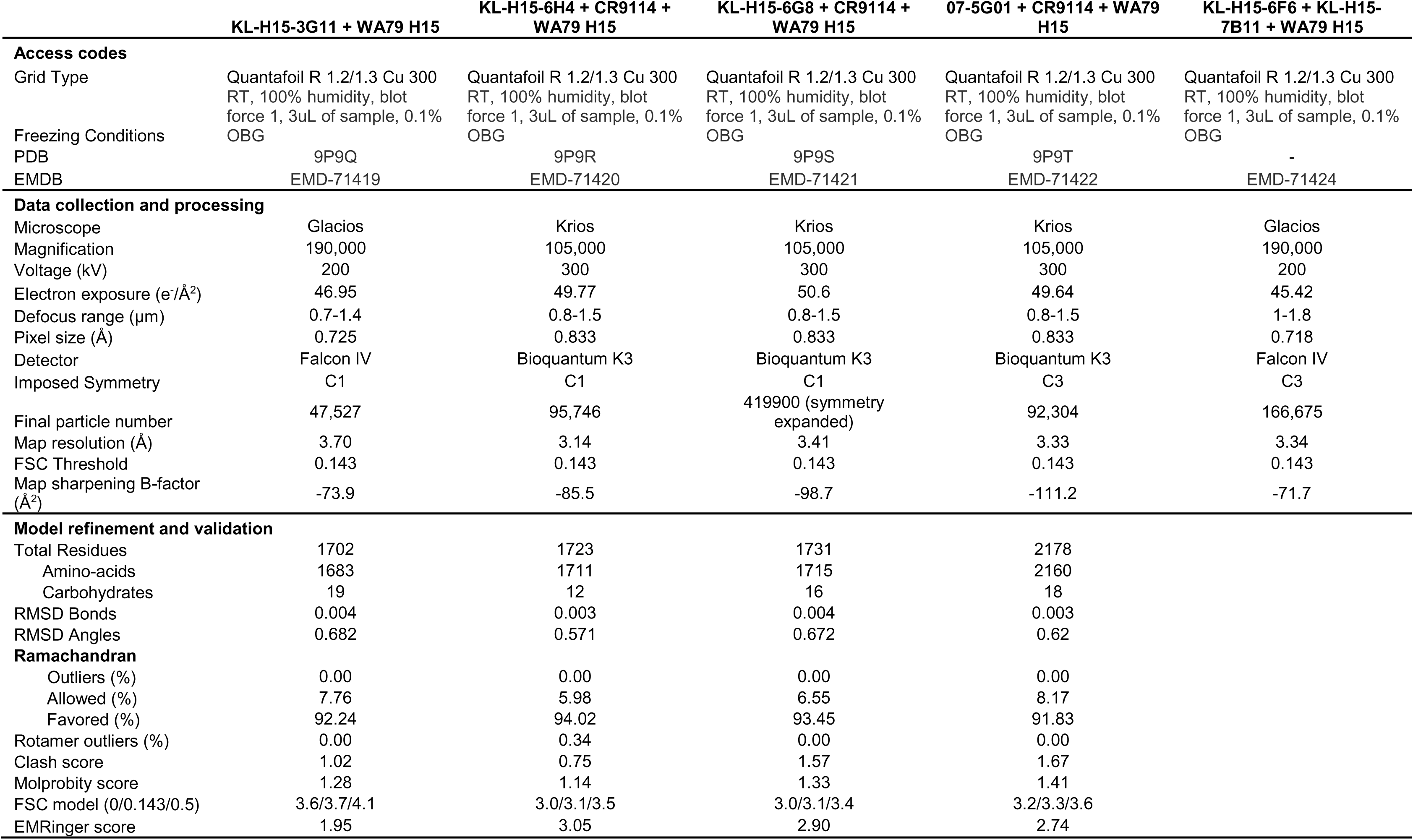
Cryo-EM data collection, processing, model refinement and validation statistics. Validation statistics were generated using Phenix 1.21.2 Comprehensive Validation Tool (Barad et al., 2015; Chen et al., 2010).

**Table S2.**
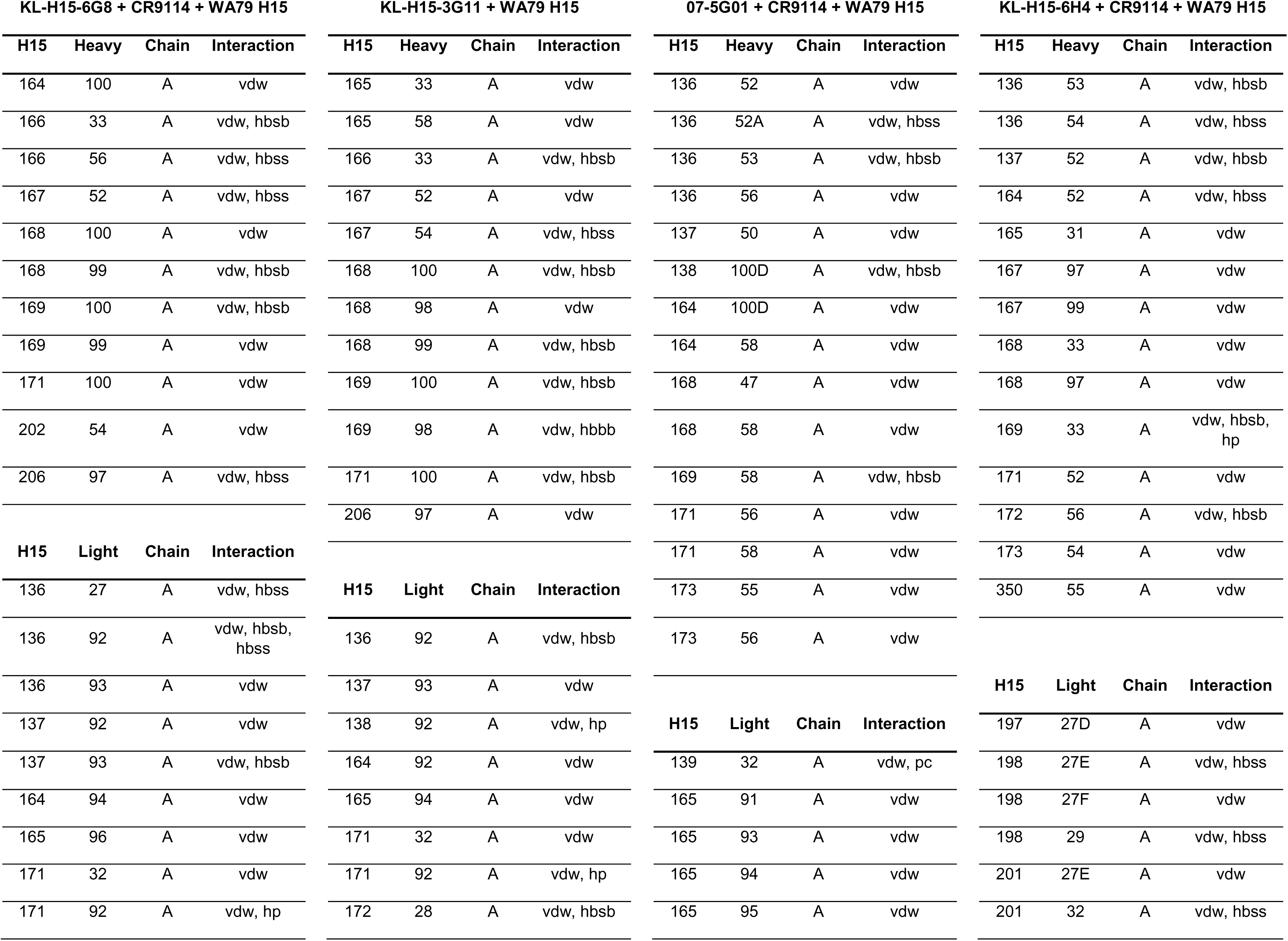
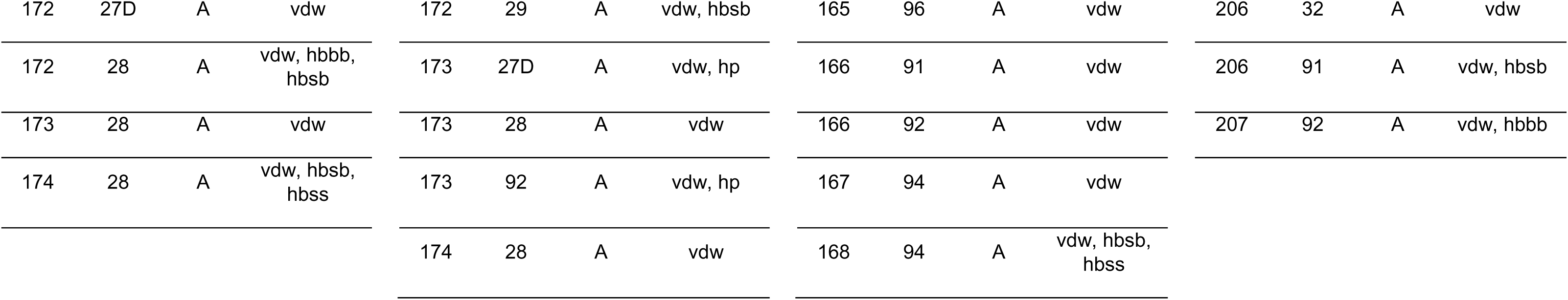
Epitope-paratope contacts predicted from cryoEM models. The GetContacts program (https://getcontacts.github.io/index.html) was used to predict residue interactions at the epitope-paratope. Interactions are abbreviated as follows: van der Waals (vdw), side-chain to backbone hydrogen bonding (hbsb), backbone to backbone hydrogen bonding (hbbb), side-chain to side-chain hydrogen bonding (hbss), and hydrophobic (hp).

**KL-H15-6G8 + CR9114 + WA79 H15**
| H15 | Heavy | Chain | Interaction |
| --- | --- | --- | --- |
| 164 | 100 | A | vdw |
| 166 | 33 | A | vdw, hbsb |
| 166 | 56 | A | vdw, hbss |
| 167 | 52 | A | vdw, hbss |
| 168 | 100 | A | vdw |
| 168 | 99 | A | vdw, hbsb |
| 169 | 100 | A | vdw, hbsb |
| 169 | 99 | A | vdw |
| 171 | 100 | A | vdw |
| 202 | 54 | A | vdw |
| 206 | 97 | A | vdw, hbss |
| H15 | Light | Chain | Interaction |
| 136 | 27 | A | vdw, hbss |
| 136 | 92 | A | vdw, hbsb, hbss |
| 136 | 93 | A | vdw |
| 137 | 92 | A | vdw |
| 137 | 93 | A | vdw, hbsb |
| 164 | 94 | A | vdw |
| 165 | 96 | A | vdw |
| 171 | 32 | A | vdw |
| 171 | 92 | A | vdw, hp |

**KL-H15-3G11 + WA79 H15**
| H15 | Heavy | Chain | Interaction |
| --- | --- | --- | --- |
| 165 | 33 | A | vdw |
| 165 | 58 | A | vdw |
| 166 | 33 | A | vdw, hbsb |
| 167 | 52 | A | vdw |
| 167 | 54 | A | vdw, hbss |
| 168 | 100 | A | vdw, hbsb |
| 168 | 98 | A | vdw |
| 168 | 99 | A | vdw, hbsb |
| 169 | 100 | A | vdw, hbsb |
| 169 | 98 | A | vdw, hbbb |
| 171 | 100 | A | vdw, hbsb |
| 206 | 97 | A | vdw |
| H15 | Light | Chain | Interaction |
| 136 | 92 | A | vdw, hbsb |
| 137 | 93 | A | vdw |
| 138 | 92 | A | vdw, hp |
| 164 | 92 | A | vdw |
| 165 | 94 | A | vdw |
| 171 | 32 | A | vdw |
| 171 | 92 | A | vdw, hp |
| 172 | 28 | A | vdw, hbsb |

**07-5G01 + CR9114 + WA79 H15**
| H15 | Heavy | Chain | Interaction |
| --- | --- | --- | --- |
| 136 | 52 | A | vdw |
| 136 | 52A | A | vdw, hbss |
| 136 | 53 | A | vdw, hbsb |
| 136 | 56 | A | vdw |
| 137 | 50 | A | vdw |
| 138 | 100D | A | vdw, hbsb |
| 164 | 100D | A | vdw |
| 164 | 58 | A | vdw |
| 168 | 47 | A | vdw |
| 168 | 58 | A | vdw |
| 169 | 58 | A | vdw, hbsb |
| 171 | 56 | A | vdw |
| 171 | 58 | A | vdw |
| 173 | 55 | A | vdw |
| 173 | 56 | A | vdw |
| H15 | Light | Chain | Interaction |
| 139 | 32 | A | vdw, pc |
| 165 | 91 | A | vdw |
| 165 | 93 | A | vdw |
| 165 | 94 | A | vdw |
| 165 | 95 | A | vdw |

**KL-H15-6H4 + CR9114 + WA79 H15**
| H15 | Heavy | Chain | Interaction |
| --- | --- | --- | --- |
| 136 | 53 | A | vdw, hbsb |
| 136 | 54 | A | vdw, hbss |
| 137 | 52 | A | vdw, hbsb |
| 164 | 52 | A | vdw, hbss |
| 165 | 31 | A | vdw |
| 167 | 97 | A | vdw |
| 167 | 99 | A | vdw |
| 168 | 33 | A | vdw |
| 168 | 97 | A | vdw |
| 169 | 33 | A | vdw, hbsb, hp |
| 171 | 52 | A | vdw |
| 172 | 56 | A | vdw, hbsb |
| 173 | 54 | A | vdw |
| 350 | 55 | A | vdw |
| H15 | Light | Chain | Interaction |
| 197 | 27D | A | vdw |
| 198 | 27E | A | vdw, hbss |
| 198 | 27F | A | vdw |
| 198 | 29 | A | vdw, hbss |
| 201 | 27E | A | vdw |
| 201 | 32 | A | vdw, hbss |

|  |  |  |  |  |  |  |  |  |  |  |  |  |  |  |  |
| --- | --- | --- | --- | --- | --- | --- | --- | --- | --- | --- | --- | --- | --- | --- | --- |
| 172 | 27D | A | vdw | 172 | 29 | A | vdw, hbsb | 165 | 96 | A | vdw | 206 | 32 | A | vdw |
| 172 | 28 | A | vdw, hbbb,<br>hbsb | 173 | 27D | A | vdw, hp | 166 | 91 | A | vdw | 206 | 91 | A | vdw, hbsb |
| 173 | 28 | A | vdw | 173 | 28 | A | vdw | 166 | 92 | A | vdw | 207 | 92 | A | vdw, hbbb |
| 174 | 28 | A | vdw, hbsb,<br>hbss | 173 | 92 | A | vdw, hp | 167 | 94 | A | vdw |  |  |  |  |
|  |  |  |  | 174 | 28 | A | vdw | 168 | 94 | A | vdw, hbsb,<br>hbss |  |  |  |  |

